# Word predictability, prosody, gesture and mouth movements in face-to-face language comprehension

**DOI:** 10.1101/2020.01.08.896712

**Authors:** Ye Zhang, Diego Frassinelli, Jyrki Tuomainen, Jeremy I Skipper, Gabriella Vigliocco

## Abstract

The ecology of human language is face-to-face interaction, comprising cues, like prosody, co-speech gestures, and mouth movements. Yet, the multimodal context is usually stripped away in experiments as dominant paradigms focus on linguistic processing only. In two studies we presented video-clips of an actress producing naturalistic passages to participants whose electroencephalographic activity was recorded. We quantified each cue and determined their effect on a well-established electroencephalographic marker of cognitive load in comprehension (N400). We found that brain responses to words were affected by informativeness of co-occurring multimodal cues, indicating that comprehension relies on linguistic and non-linguistic cues. Moreover, brain responses were affected by interactions between the multimodal cues, indicating that the impact of each cue dynamically changes based on the informativeness of other available cues. Thus, results show that multimodal cues are integral to comprehension, hence, our theories must move beyond the limited focus on speech and linguistic processing.

## Introduction

Language originated, is learned and most often used in face-to-face settings. In these contexts, linguistic information is accompanied by other multimodal ‘non-linguistic’ cues like speech intonation (prosody), hand gestures and mouth movements. Behavioural, neuroimaging and electrophysiological research has shown that these cues can individually improve speech perception and language comprehension in experimental studies in which the other cues are carefully controlled or absent^1–4^. However, in real-world scenarios these cues often co-occur. Yet, most theoretical accounts of language comprehension are grounded in studies focusing only on a single (usually ‘linguistic’) cue. Taken together, this limits the ecological validity of existing studies and theories.

There is increasing evidence that prediction plays a general role in brain functioning, including (audiovisual) speech perception and language comprehension^5–7^. Predictions matter in speech and language because they might provide a constraint on the interpretation of ambiguities in the acoustic signal, at word and higher linguistic levels. Many previous studies have demonstrated the role of prior linguistic material (e.g., prior discourse) in developing predictions for upcoming sounds or words. Co-occurring multimodal cues such as prosody, gestures and mouth movements can modulate such predictions^8,9^. However, because most often studies do not include these ‘non-linguistic’ cues, the mechanisms underscoring natural language comprehension in the brain are unclear. In particular, there are - at least - two key questions that we need to answer at first in order to develop comprehensive theories. First, we need to understand to what extent the processing of multimodal cues is central in natural language processing (e.g., whether a cue is only used when the speech is ambiguous, or in experimental tasks that force attention to it). Answering this question is necessary in order to properly frame theories of natural language processing because, if some multimodal cues (e.g., gesture or prosody) *always* contribute to processing, this would imply that our current speech-only focus is too narrow, if not misleading. Second, we need to understand the dynamics of online multimodal comprehension. To provide mechanistic accounts of language comprehension, it is necessary to establish how the weight of a certain cue dynamically changes depending upon the presence and informativeness of other cues (e.g., whether meaningful hand gestures are weighted more when prior linguistic context is less informative and/or when mouth movements are less informative).

### The impact of prosody, gesture and mouth movements on processing: the state-of-the-art

Accentuation (i.e., prosodic stress characterized as higher pitch that makes words acoustically prominent) marks new information^10^. Many behavioural studies have revealed that comprehension is facilitated with appropriate accentuation (new information is accentuated, and old information de-accentuated^11,12^). Incongruence between the presence of prosodic accentuation and newness of information increases cognitive load during language comprehension. It induces an increased activation in the left inferior frontal gyrus, indicating increased difficulty in phonological and semantic processing^13^. In electrophysiological (EEG) studies, this mismatch elicits a more negative N400 than appropriate accentuation^14–18^. The N400 is an event-related-potential (ERP) peaking negatively around 400ms after word presentation around central-parietal areas^19^, that has been argued to index cognitive load and prediction in language comprehension^2^.

Meaningful co-speech gestures have been shown to improve comprehension by providing additional semantic information about upcoming words^20^. In line with behavioural studies, EEG studies have shown that activating the less predictable meaning of the homonymous word “ball” using a “dancing” gesture, reduces the N400 response to a later mention of “dance”^21–23^. Incongruence between meaningful gestures and linguistic context triggers more negative N400 compared with congruent gestures,indicating that meaningful gestures can constrain predictions for upcoming words based on previous linguistic context^24–27^. Processing of co-speech meaningful gestures has been linked to activation in posterior middle-temporal and inferior frontal regions, which are associated with meaning processing across linguistic and non-linguistic materials^9,28–31^. Moreover, the presence of meaningful gestures has been shown to result in a significant reduction in cortical activity in auditory language regions (namely posterior superior temporal regions), a hallmark of prediction^32^.

Fewer studies have investigated beat gestures (meaningless gestures time-locked to the speech rhythm)^33^. Some argued that beats enhance saliency of associated speech in a similar manner as prosodic accentuation^34^, and activate the same regions as prosody in auditory cortex^35^. Two studies reported that beat gestures induce less negative N400, similar to prosodic accentuation^36,37^. Other EEG studies, however, reported that beat gestures modulated brain responses in a later window (around 600ms^38,39^).

Finally, many previous studies focused on the sensory-motor mechanism underscoring the use of mouth movements in speech, indicating that mouth movements facilitate the perception of auditory signal^40,41^, modulate early sensory electrophysiological signals (N1-P2)^42,43^, and activate auditory cortices.^44,45^ Less is known about whether the informativeness of mouth movements affects word predictability. Some behavioural work suggest that mouth movement may facilitate lexical access^46–48^ and meaning comprehension^49^. Other behavioural and fMRI studies show that the facilitatory effect of mouth movements persists even with the presence of meaningful gestures^9,30,50^. Two electrophysiological studies, however, reported conflicting findings. While Brunellière and colleagues linked more informative mouth movements to more negative N400 amplitude^51^, generally indicating increased processing difficulty, Hernández-Gutiérrez and colleagues failed to find any N400 effect associated with mouth movements^52^.

Thus, previous studies indicate that at least when taken one by one, multimodal non-linguistic cues interact with speech in modulating the predictability of upcoming words. Because of the challenges of using naturalistic stimuli, often in these studies the investigated cues are carefully manipulated while all the other possible interacting cues are either eliminated or kept constant^53^. Thus, for example, prosody is normalised and auditory (rather than audiovisual) presentation is used when studying speech^54^; only the mouth, rather than the whole body is shown when studying audiovisual speech perception^51^; and the face is hidden when studying gestures^21^. In this way, the materials and tasks often do not reflect the conditions in which the brain processes language in real-world face-to-face contexts: it is simply impossible not to see a person’s gestures while they speak, or their mouth movements while we see their gestures. Furthermore, the standard reductionist paradigm breaks the natural and possibly predictive correlation among cues with unknown consequences on processing^55,56^. The disruption of the relative reliability of cues can affect whether and how much the brain relies on a given cue^57,58^.

### The present study

Using a design that preserves ecological validity, we address two key questions about face-to-face multimodal communication: (1) to what extent is the processing of multimodal (‘non-linguistic’) cues central to natural language processing? and (2) what are the dynamics of online multimodal comprehension? We address these questions investigating whether the presence of any multimodal cue (and their combination) modulates predictions - based on prior discourse - for upcoming words. We carried out two experiments (an original study and a replication with different materials) using materials that preserve the natural correlation across cues. In the first experiment, we asked thirty-six (31 included, mean age=27, 17 women) native English speakers to watch 100 videos in which an actress produced short passages (taken from a corpus representing an extensive variety of British English natural language usage)^59^ with natural prosody, co-speech gestures and mouth movements. We replicated this study asking another 20 native English speaking participants (20 included, mean age=25, 15 women) to watch 79 videos in which the same actress produced passages selected from TV transcripts. In both experiments, some videos (Exp1: n=35; Exp2: n=40) were followed by yes/no questions about the content of the video to ensure participants’ attention (and to acquire behavioural responses in Experiment 1, see Supplementary Materials). Participants were instructed to watch the videos carefully and to answer the questions as quickly and accurately as possible. We measured the electrophysiological responses to each word and assessed how each cue and their interactions modulate N400 responses. We use the N400 as a biological marker of cognitive load, associated with word predictability^60^. Crucially as discussed above, prosody, gestures and mouth movements have all been shown to modulate N400 responses to words, indicating that the time window in which we observe this event-related potential is suitable to detect any effect of the multimodal cues on word processing.

In both Experiments, word-predictability was computed for each content word in the passages using semantic surprisal, a measure indicating the probability of encountering a specific word following a given linguistic context^60,61^. Prosody (prosodic accentuation) was quantified in terms of the mean fundamental frequency (F0) of each word^62^; gestures (meaningful gestures and beats) were coded as present/absent for each word and finally, mouth movements associated to each word were quantified in terms of their informativeness (i.e., how easy it is to guess the word just by looking at the corresponding mouth movements). Quantification of the different cues word-by-word allows us to address how their dynamic change impacts electrophysiological responses. Figure 1 gives an example of an annotated sentence.

**Figure 1.**
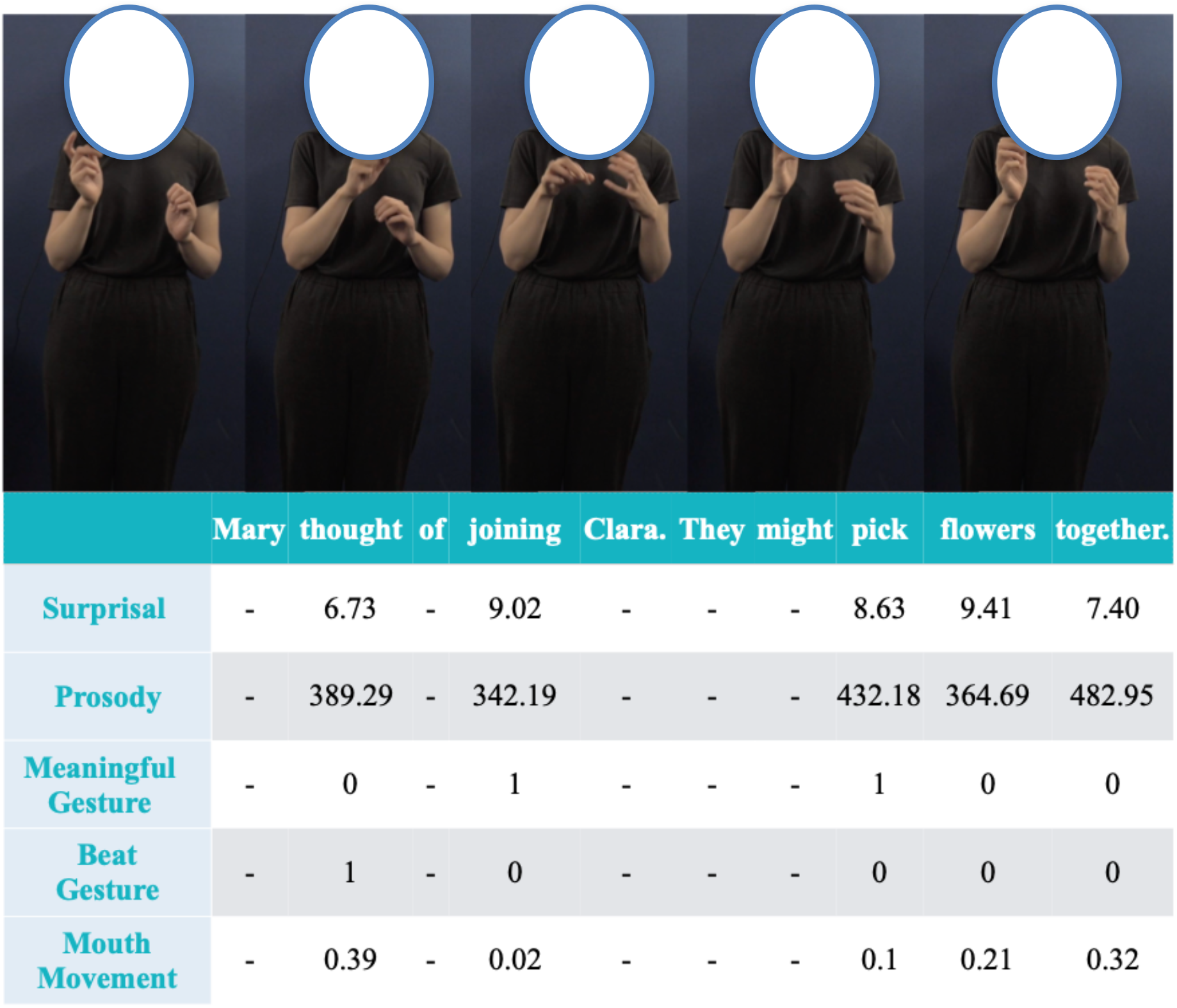
Example of stimuli and annotations. Annotation was carried out for content words only. Each frame corresponds to an image during each such word.

To assess whether the processing of these multimodal cues is central in natural language processing, we measured whether the presence/informativeness of multimodal non-linguistic cues modulates the impact of linguistic predictability, indexed using the N400. Previous work suggests that prosody and meaningful gestures will reduce the N400 amplitude as both provide meaningful information that makes words that would be less predictable in the prior linguistic context easier to process. Here, we go beyond this by asking whether the same pattern will hold when it is not just one, but multiple cues contributing to the process. Second, we evaluate the dynamic nature of multimodal cue processing by analyzing the interaction between cues. If the weight of a certain cue dynamically changes depending upon the context, then its impact on word predictability should change as a function of other cues.

## Results

### EEG Analyses

#### Time Window Sensitive to Linguistic Context

Experiment 1’s data was first used to establish the time-window in which processing is affected by linguistic predictability. No previous study has investigated the effect of surprisal in audiovisual multimodal communication. Therefore, rather than making a priori assumptions about the specific event-related response we should observe and the time window in which we would observe it, we carried out a hierarchical LInear MOdeling (LIMO toolbox^63^) to establish the EEG component sensitive to surprisal. While traditional ERP analysis compares different conditions and thus may require dichotomization of the predictor variable^64^, this regression based ERP analysis linearly decomposes the ERP signal into timeseries of beta coefficient waveforms elicited by continuous variables. Significant differences between the beta coefficient waveforms and zero (or a flat line, indicating that the variable does not affect EEG signal) represent the existence of an effect^65,66^. We found that the beta values of central-parietal electrodes were significantly different from zero in the 300-600ms time window across electrodes (Figure 2). Words with higher surprisal, elicited more negative signals or larger N400 amplitudes. No other time window was significantly sensitive to surprisal. As a result, we focused on the 300-600ms interval in our subsequent analyses in both studies (analysis of Experiment 2 yielded approximately the same window; see S.M.).

**Figure 2.**
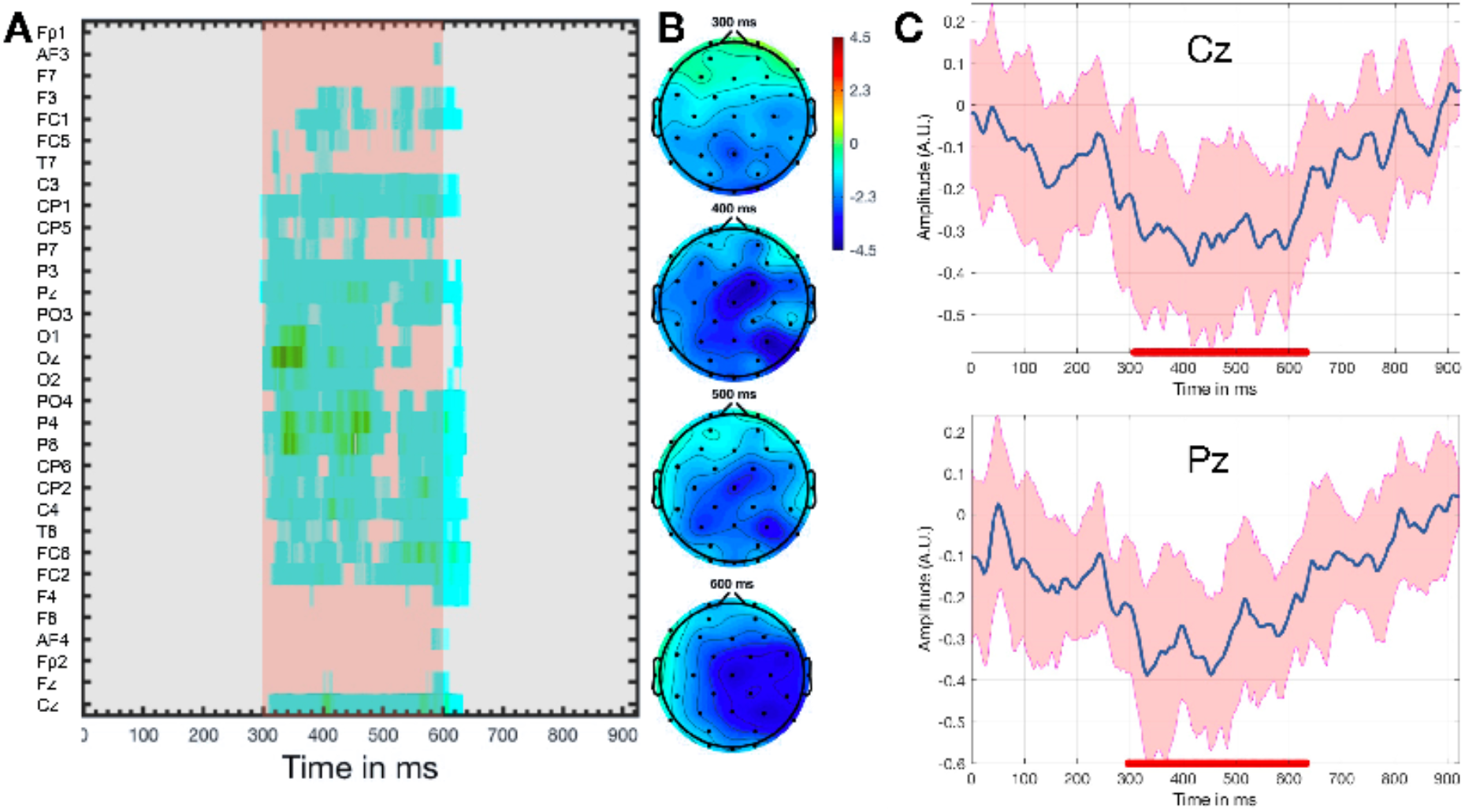
Hierarchical linear modelling results showing the ERP sensitive to surprisal (one-sample t-test P<0.05, cluster-corrected). (A) Time window (300-600ms) showing increased significant negativity associated with surprisal (in pink). Grey areas are not statistically significant. (B) Topographic maps illustrating the scalp distribution for the 300-600 time window. Deeper blue area indicates more negative beta values. (C) Averaged beta plot for electrode Cz and Pz illustrating that beta values for surprisal were significantly negative compared with 0 (flat waveform) in 300-600ms. The blue line indicates the average beta value, while red indicates the confidence interval. The red line underlying the figures indicates the significant time window. Cz and Pz are chosen here because they are most often used to depict N400 effects (that are maximal at central-parietal locations^8^)

### Multimodal cues are central to language processing: Modulation of word predictability by the cues

After determining the correct time window, we performed a linear mixed effect analysis (LMER) on the averaged ERP amplitude in the 300-600ms window. LMER was used due to the advantage in accommodating both categorical and continuous variables, thus increasing statistical power^67^. In the following studies, we adopted the most complex fixed and random model structure allowed by the data as suggested by (Ref, Keep it Max). Moreover, LMER can account for both by participant and item (word lemmas) variance and can better accommodate unbalanced designs; such properties make it suitable for EEG studies investigating naturalistic language processing^60,68^.

Mean ERP in the 300-600ms time window was used as the dependent variable. The independent variables are: surprisal, mean F0, meaningful gestures, beat gestures, mouth informativeness and all up to three-way interactions between surprisal and any other two cues, alongside other control variables (see Methods). We further included word lemma and participant as random intercepts. The highest interactions (all three-way interactions) were also included as random slopes for participant^69^, and surprisal as random slope for lemma. The random slope of surprisal for lemma was removed in the replication study due to convergence issues. Full model results and ERP plots are reported in S.M.

We focus first on the main effects of the multimodal cues and their interaction with surprisal. In this way, we can assess the exact contribution of the multimodal cues to the purely linguistic processing of a word described by surprisal.

As shown in Figure 3 (panel A), we found a main effect of prosody (mean F0) (Exp1: β=0.010, p<.001; Exp2: β=0.014, p<.001). Words produced with higher mean pitch showed less negative EEG, or smaller N400 amplitude, compared with words produced with lower pitch. Figure 3, panel B reports the interaction between surprisal and mean F0 (Exp1: β=0.017, p<.001; Exp2: β=0.012, p<.001). The larger N400 amplitude associated with high surprisal words was modulated by pitch. High surprisal words elicited a larger reduction of N400 amplitude when the pitch was higher, in comparison to low surprisal words.

**Figure 3.**
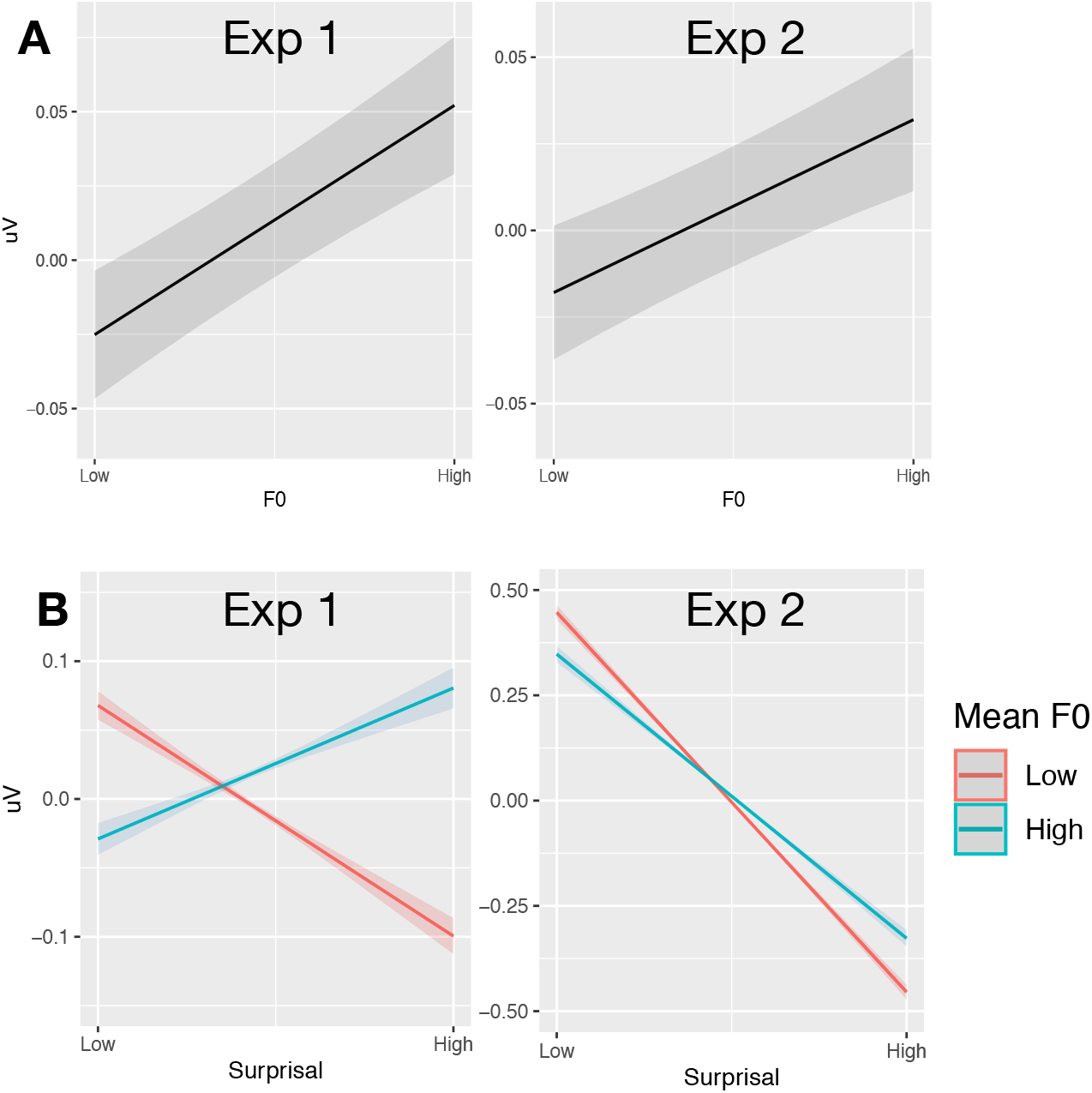
Prosodic Accentuation (mean F0) modulation of N400 amplitude. (A) Main effect of Prosodic Accentuation. (B) Interaction between Prosodic Accentuation and Surprisal. Plots depict the predicted value of the mean amplitude of the ERP within 300-600ms (grey areas = confidence intervals).

We found a similar main effect of meaningful gestures (Figure 4). Words accompanied by a meaningful gesture showed a significantly less negative N400 (Exp1: β=0.006, p<.001; Exp2: β=0.007, p<.001). There was also a significant interaction between surprisal and meaningful gesture, indicating that for high surprisal words, the presence of a meaningful gesture makes the N400 less negative (Exp1: β=0.008, p<.001; Exp2: β=0.011, p<.001).

**Figure 4.**
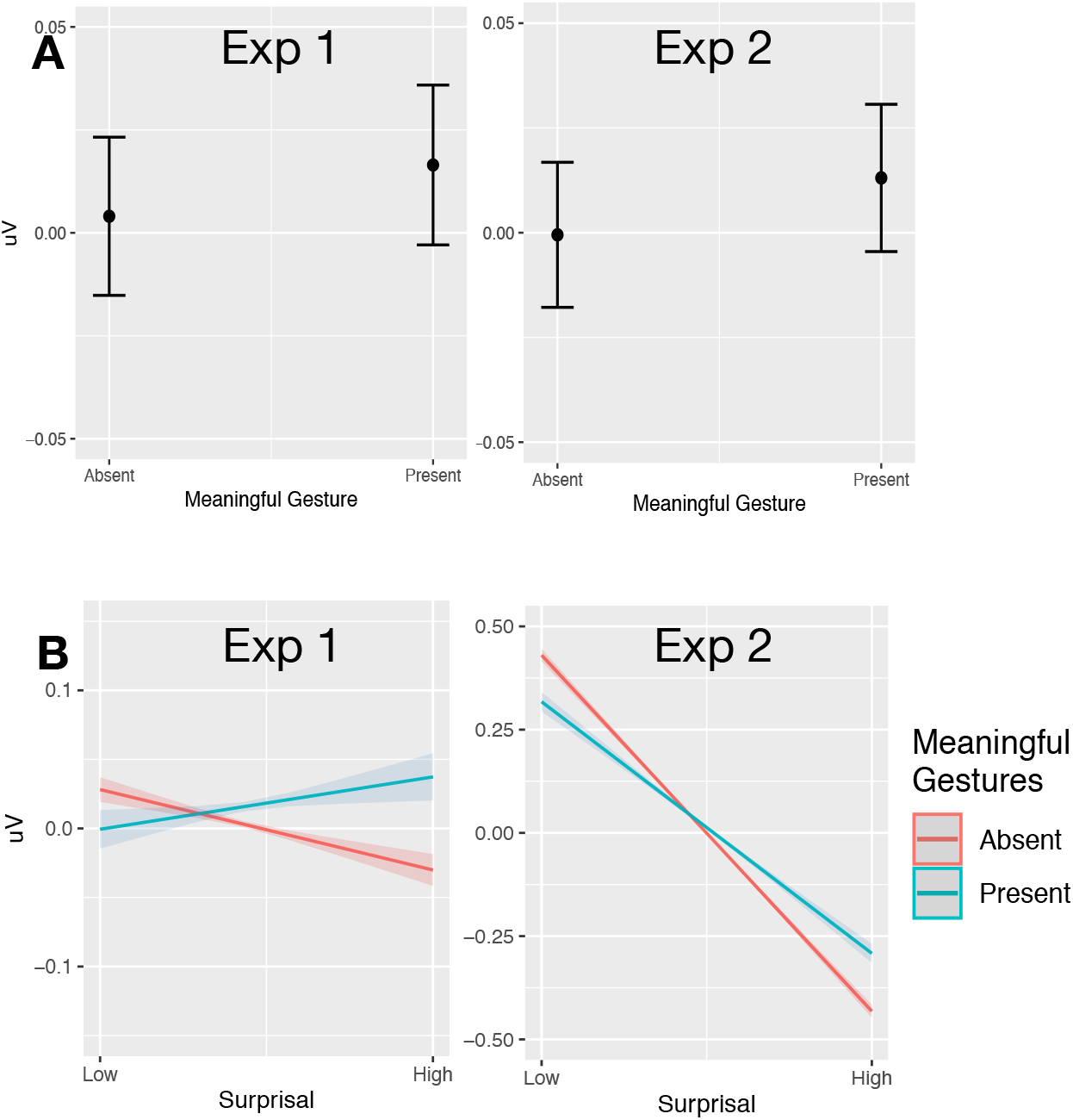
Meaningful gesture modulation of N400 amplitude. (A) Main effect of meaningful gestures. (B) Interaction between Meaningful Gestures and Surprisal. Conventions are the same as in Figure 3.

In contrast to meaningful gestures, beat gestures showed a different pattern (Figure 5). We found a significant main effect of beat gestures (Exp1: β=-0.005, p=.001; Exp2: β=-0.006, p=.001), suggesting that words accompanied by beat gestures elicited a more negative N400. Moreover, the high surprisal words accompanied by beat gestures showed a further significant increase in negativity compared with low surprisal words (Exp1: β=-0.012, p<.001; Exp2: β=-0.010, p<.001).

**Figure 5.**
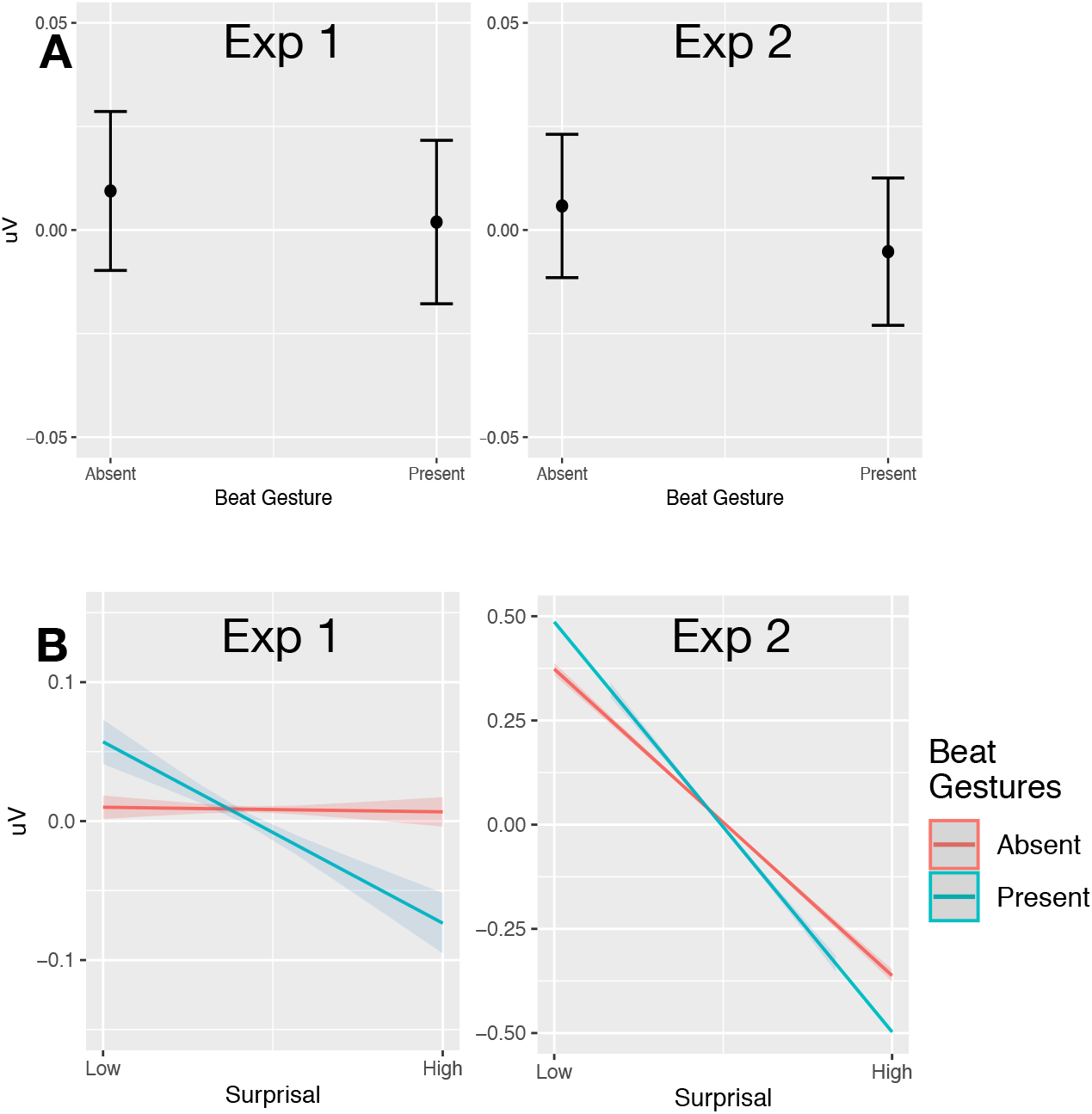
Beat gesture modulation of N400 amplitude. Main effect of beat gestures. (B) Interaction between Beat Gestures and Surprisal. Conventions are the same as in Figure 3.

### Dynamics of Multimodal Cue Processing: Interactions among Multimodal Cues

We found a number of significant interactions between multimodal cues (Figure 6). First, there was an interaction between mean F0 and meaningful gesture (Exp1: β=0.004, p<.001; Exp2: β=0.005, p<.001). Words with meaningful gestures showed even less negative amplitude of N400 with increased mean F0. There is also a significant interaction between mouth informativeness and meaningful gesture (Exp1: β=0.004, p=0.002; Exp2: β=0.007, p<. 001) and between mouth informativeness and beat gesture (Exp1: β=0.012, p<0.001; Exp2: β=0.004, p=0.001). Words with more informative mouth movement showed less negative N400 with the presence of both meaning fuland beat gestures.

**Figure 6.**
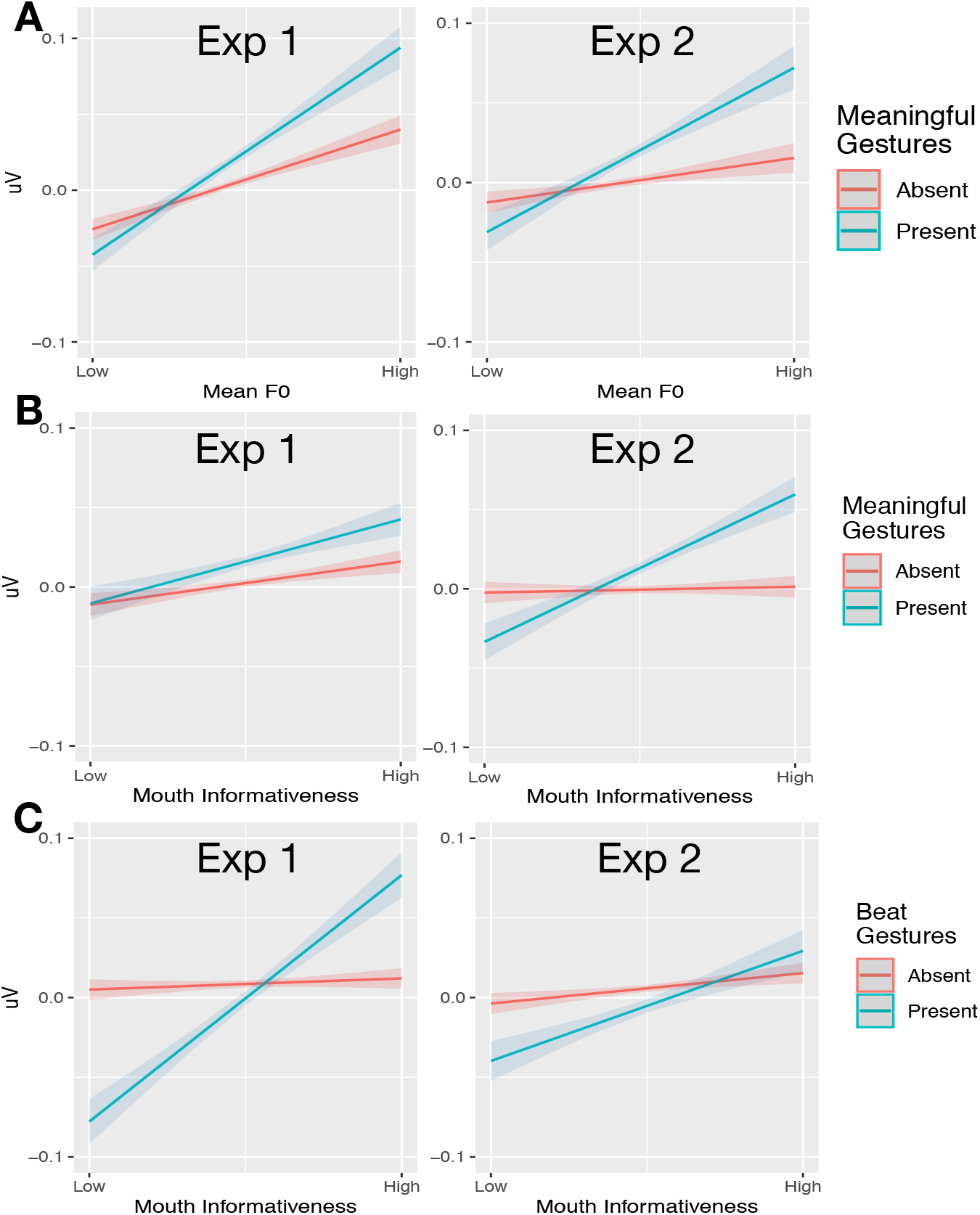
Interactions between multimodal cues. (A) Interaction between Prosodic Accentuation and Meaningful Gestures. (B) Interaction between Meaningful Gestures and Mouth Informativeness. (C) Interaction Beat Gestures and Mouth Informativeness. Conventions are the same as in Figure 3.

## Discussion

The present study investigated for the first time the electrophysiological correlates of real-world multimodal language comprehension tracking the on-line processing as indexed by N400 amplitude. Unlike most previous studies, our stimuli did not break the naturally occurring correlations among multimodal cues. First, we confirmed the N400 as a biomarker of prediction during naturalistic audiovisual language comprehension: high surprisal words elicited a more negative N400 between 300 and 600 ms post-stimulus, strongest in the central-posterior electrodes.

More broadly, our study provides the first comprehensive picture of how the brain dynamically weights audiovisual cues in real-world language comprehension and it provides answers to the two key questions we presented in the introduction. First, we asked whether the processing of these multimodal cues is central to natural language processing. We found that this is the case as indexed by the fact that each cue (except mouth informativeness) had a general effect and crucially modulated linguistic-based surprisal. Prosodic accentuation and meaningful gestures reduced the N400 amplitude overall, especially for high surprisal words. In contrast, the presence of beat gestures increased the N400 amplitude overall, especially for high surprisal words. Mouth movements did not modulate surprisal independently, but participated in complex interactions involving other cues. Thus, our results clearly show that language comprehension in its natural ecology, as face-to-face communication, involves more than just speech: the predictability of words based on linguistic context is *always* modulated by the multimodal cues thus forcing a reconsideration of theoretical claims strictly based on speech or linguistic processing only^8^.

Second, we addressed the dynamic nature of multimodal cue processing. We found that the weight given to each cue at any given time depends on which other cue is present, as indexed by interactions between cues. We found that first, the facilitatory effect of meaningful gestures was enhanced when the word had higher pitch. Second, the facilitatory effect of mouth informativeness was enhanced when either meaningful or beat gestures were present. Thus, these results show that investigating one cue at the time does not provide the full picture precluding the development of a mechanistic understanding of multimodal processing.

### Prosody, gesture and mouth movements contribution to linguistic processing: beyond the state of the art

By using more naturalistic stimuli that do not artificially isolate a single cue at the time, our results provide clarifications on previous conflicting results and offer additional insight into cue interactions.

Prosodic accentuation has been considered to mark ‘newness’^1^, as speakers are more likely to stress a word if it conveys new information^10^. Previous electrophysiological studies have shown that un-accented new words elicit more negative N400^14–18,70^. Our findings complement previous work in showing that in multimodal contexts, the presence of accentuation for less predictable words reduces the amplitude of the N400, suggesting that prosodic accentuation can enhance expectation for lower probability continuations, in line with earlier behavioural works^11,12^. We found that meaningful gestures support processing, especially for high surprisal words. This result is in line with studies that showed N400 reduction for the subordinate meaning of ambiguous words (e.g. “ball” meaning dancing party) in the presence of a corresponding gesture^21–23^, and previous work suggesting that words produced with incongruent gestures induce a larger N400 (see review in Özyürek, 2014^4^). Our results show that meaningful gestures play a more general role in face-to-face communication: meaningful gestures are always supporting word processing, not just in cases in which processing is especially difficult (due to incongruence or ambiguity).

Crucially, *meaningful* gestures, but not beat gestures, decrease the cognitive load associated with word processing. High surprisal words accompanied by beat gestures elicited an even larger N400 effect. This effect might be accounted for in terms of beats enhancing the saliency of a specific word^34^, and highlighting its lack of fit into the previous context. Alternatively, it is possible that listeners try to extract meaning from all gestures and integrate it with speech by default, and since beats are not meaningful, integration fails, inducing processing difficulties. Importantly, the dissociation between meaningful and beat gestures further allows us to exclude the possibility that the N400 reduction observed (for meaningful gestures and for prosody) comes about because these multimodal cues share processing resources with speech processing, letting less predictable words go unnoticed.

Previous studies failed to find the same effects of beat gestures within the N400 time window^36,37^. However, these studies used artificial beat gestures (one single stroke was aligned with the target word for each sentence), which are different from naturally occurring beat gestures (see also^71^). Alternatively, it could be the case that the lack of any meaningful gestures in their study could have discouraged listeners from paying attention to gestures. Shifts in the weight attributed to different multimodal cues depending upon the specific task used are documented in the literature^21,57,58^ and highlight the importance of using ecologically valid paradigms.

Based on studies investigating single cues, it has been suggested that beats and prosodic accentuation serve the same function in communication, namely, they both make individual words more prominent and therefore attract attention to them^34^. However, our results provide evidence against such a claim as their electrophysiological correlates dissociated: beat gestures elicited more negative N400 especially for high surprisal words in line with the account above, while prosodic accentuation elicited less negative N400, especially for high surprisal words (see also^36^).

We did not find a reliable effect of mouth informativeness as the main effect or in interaction with surprisal in the N400 time-window. Mouth movements have long been recognized to facilitate speech perception especially in noisy situations^60^; moreover, audiovisual communication showed a reduction in the N1/P2 amplitude compared to the audio-only condition, indicating easier sensory-level processing^42,43^. However, in our study we focused on 300-600ms after word onset in order to capture the effect of surprisal and we did not consider earlier (100-300ms) time windows. Two previous studies have investigated the impact of mouth movements within the N400 time window. Hernández-Gutiérrez and colleagues did not find any N400 difference between audiovisual and audio-only speech^52^; while Brunellière and colleagues found an increase in N400 amplitude for more informative mouth movements^51^. Further research is necessary to clarify these discrepancies, however, our results suggest that mouth informativeness can affect processing in the N400 time window but only in combination with other cues when presented in a multimodal context.

Finally, our results further extend the previous literature by showing how cues interact. We found that meaningful gestures and prosodic modulation interact. Kristensen and colleagues argued that prosodic accentuation engages a domain general attention network^72^. Thus, accentuation may draw attention to other cues which consequently would be weighted more heavily. However, while this account is plausible for meaningful gestures, it does not explain why we did not observe a similar enhancement for beat gestures. Alternatively (or additionally) as argued by Holler and Levinson, listeners are attuned to natural correlations among the cues (e.g., high pitch correlates to larger mouth movements and increased gesture size) and would use cue-bundles for prediction^8^.

Moreover, we found that the effect of mouth informativeness is enhanced whenever other visual cues like meaningful gestures or beats are present. This may happen because mouth movements would fall within the focus of visual attention more easily if attention is already drawn to gestures (it is known that listeners look at the chin when processing sign language and gestures^73^).

### Toward a neurobiology of natural communication

Our result calls for a new neurobiological model of natural language use that accounts for the effects of multimodal cues on language comprehension, as well as the interactions within multiple multimodal cues. In probabilistic-based predictive accounts, the N400 is taken as an index of the processing demands associated with low predictability words^2^. It has been argued that prior to the bottom-up information, a comprehender holds a distribution of probabilistic hypotheses of the upcoming input constructed by combining his/her probabilistic knowledge of events with contextual information. This distribution, solely based on linguistic input, is updated with new information, and consequently becomes the new prior distribution for the next event. Thus, the N400 is linked to the process of updating the distribution of hypotheses: smaller N400 is associated with more accurate prior distributions/ predictions^2^. Our work shows that these mechanisms do not operate only on linguistic information, but crucially, they weigh in ‘non-linguistic’ multimodal cues.

In terms of neuroanatomical models, those in which language comprehension is considered in context and associated with many interconnected networks distributed throughout the whole brain^55,56^ can, in principle, accommodate the results reported here. For example, in the Natural Organization of Language and Brain (NOLB) model, each multimodal cue is proposed to be processed in different but partially overlapping sub-networks^56^. Indeed, different sub-networks have been associated with gestures and mouth movements, with a ‘gesture network’ and ‘mouth network’ weighted differently in different listening contexts^9,30^. These distributed sub-networks are assumed to provide constraints on possible interpretations of the acoustic signal, thus enabling fast and accurate comprehension^30^. Our finding of multiple interactions between cues is compatible with this view, thus suggesting that multimodal prediction processes are dynamic, re-weighting each cue based on the status of other cues.

### Conclusions

To conclude, our study investigated language processing in its naturalistic multimodal environment for the first time, and provided novel and robust evidence that, first, multimodal ‘non-linguistic’ cues have a central role in processing as they always modulate predictions on what is going to be said next; second, they dynamically interact among one another and with linguistic cues to construct these predictions. More generally, our study provides a more ecologically valid way to understand the neurobiology of language, in which multimodal cues are dynamically orchestrated.

## Methods

### Participants

#### Experiment 1

Thirty-six native English speakers with normal hearing and normal or corrected to normal vision were paid £7.5 per hour to participate in the study after giving written consent. Five participants were excluded, three due to technical issues, one for falling asleep, and one for excessive muscle noise, leaving thirty-one participants.

#### Experiment 2

Twenty native English speakers with the same criteria above participated in the replication study after giving written consent.

All methods were carried out in accordance with relevant guidelines and regulations, and all experimental protocols were approved by the University College London ethics committee.

### Material

#### Experiment 1

Two-hundred and forty-six naturalistic sentence pairs (two consecutive sentences) were extracted from the British National Corpus (BNC, University of Oxford, 2007)^59^. Sentences were selected in a semi-random fashion with the only constraints that the second sentence had to be at least five words long, and contain at least one verb that could be easily gestured (e.g. “turn the pages”). If necessary, we edited slightly the first sentence to facilitate readability and resolved all ambiguities (e.g. proper nouns without a clear reference were changed into pronouns), while the second sentence was kept unmodified. Twelve native English speakers were paid £2 each to evaluate the sentence pairs for grammaticality, meaningfulness and gesturability on a 1-5 likert scale. We selected 103 sentence pairs that had averaged gesturability > 2 (and SD < 2.5); and had no grammatical errors or semantic anomalies. Three sentence pairs were used for practice, and 100 were used as stimuli (Mean gesturability=2.67, SD=0.58).

A native British English-speaking actress produced the 103 sentence pairs. She stood in front of a dark-blue background, wearing black T-shirt and trousers to keep her arms and hands visible, and did not wear glasses to keep her face visible. She was instructed to read out the sentences presented behind the camera at a natural speed, with natural prosody and facial expressions. Each sentence pair was recorded with and without gestures. For videos with gestures, the actress was instructed to gesture as she naturally would. For videos without gestures, she was asked to stand still keeping her arms along her body. In the analyses below we compare words in the “with-gestures” condition to the same words in the “without gestures” condition. This was done because words likely to be accompanied by meaningful gestures (e.g., *combing*) are semantically very different from words that are less likely to be accompanied by gestures (e.g., *pleasing*) thus making the comparison between words produced with gestures and words produced without gestures in the same (with gesture) condition less clear. The actress has given informed consent for publication of identifying information and images in an online open-access publication.

#### Experiment 2

To better approximate the real-life language use, we chose spoken passages from TV scripts from the BBC script library (https://www.bbc.co.uk/writersroom/scripts) for the replication study. Forty-two English speakers recruited from Prolific (https://www.prolific.co/) were paid £6 per hour to rate the chosen passages on gesturability (on a Likert scale from 0 to 5; defined in the experiment as how easily gestures could be made when uttering the sentence) as well as whether the sentence was meaningful and grammatically acceptable (with “Yes” or “No”) in an online task developed using Gorilla (https://gorilla.sc/). 83 passages were included, all had 1) the mean gesturability score above 2; 2) more than 70% of the participants indicated it was grammatical; and 3) more than 70% of the participants indicated it was meaningful. Four passages were used for practice, and 79 were used as stimuli (Mean gesturability=2.89, SD=0.47). The same actress produced the passages with or without gesture under the same instructions.

### Quantification of Cues

For both studies, the onset and offset of each word were automatically detected using a word-phoneme aligner based on a Hidden Markov Model^74^. The timing was then checked and corrected manually if needed. The mean word duration was 440ms (SD=376ms) for Experiment 1, and 508ms (SD=306ms) for Experiment 2. Next, for each content word (i.e., nouns, adjectives, verbs and adverbs) we quantified the informativeness of different multimodal cues. We did not quantify measures of informativeness for function words (i.e., articles, pronouns, auxiliary verbs and prepositions) because Frank and colleagues failed to show any effect of the predictability (measured as surprisal) for such words^60^. <u>Linguistic predictability</u> was measured using surprisal (Experiment 1: Mean surprisal=7.92, SD=2.10; Experiment 2: Mean surprisal=8.17, SD=1.92), defined as the negative log likelihood of the probability of a word to follow a sequence of other words^75^. Previous work has shown that surprisal provides a good measure of predictability, especially for low predictability words^61^ and predicts reading times^61,76^ and N400 amplitude^60^. Here, surprisal was generated using a bigram language model trained on the lemmatized version of the first slice (~31-million tokens) of the ENCOW14-AX corpus, an English web corpus^77^. We chose a bigram model to reduce data sparsity and, consequently, increase the robustness of our surprisal measures. Moreover, Frank and colleagues showed that bigram models perform equally well, if not Electrophysiology of multimodal comprehension 26 better than more complex models - trigram, recurrent neural networks (RNN) and probabilistic phrase-structure grammar (PSG) - in fitting N400 data^60^. Once trained, the bigram model was used to calculate the surprisal of each word in the sentence pairs based on previous content words in the two sentences as follow::

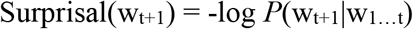

where w_t+1_ indicates the current word, and w_1…t_ stands for previous content words in the two sentences. For experiment 1, we also developed models in which the number of content words used in computing surprisal was varied. Given the minor differences observed, we decided to include all previous content words in the stimuli (from the first word to the word preceding the target word, in the supplementary material (section 1), we show results for different window sizes to justify this choice). Following this decision, Experiment 2 used all previous content words to calculate surprisal.

#### Prosodic information

for each word was quantified as its mean F0 (Experiment 1: mean F0=298Hz, SD=84Hz; Experiment 2: Mean F0=288Hz, SD=88Hz). In Experiment 1, we automatically extracted mean F0, maximum F0, minimum F0, mean intensity and F0 change per word using Praat (version 6.0.29, http://www.praat.org/)^78^. A comparison of results obtained with these different pitch measurements showed that they are similar (see supplementary material), therefore we chose to report results for mean F0 because it has been used most often as a measure of prosody in previous work^62^. Therefore, experiment 2 only extracted mean F0 following the same procedure.

#### Gestures

were coded as meaningful gestures or beats by expert coders (two in Experiment 1, three in Experiment 2). Meaningful gestures (Experiment 1: N=359; Experiment 2: N=458) comprised iconic gestures (e.g. drawing movements for the word “drawing”) and deictic gestures (e.g. pointing to the hair for “hair”). Beat gestures (Experiment1: N=229; Experiment 2: N=340) comprised rhythmic movements of the hands without clear meaning^33^, but regarded to enhance the salience of the speech^34^. Coders used ELAN (version 5.0.0, https://tla.mpi.nl/tools/tla-tools/elan/)^79^ to annotate each gesture. Gestures were annotated as iconic, deictic, beats, metaphoric or pragmatic. The phases of each gesture were coded as preparation, stroke, hold and retraction for left and right hand respectively. The lexical affiliate of each meaningful gesture was also annotated. Two variables, meaningful gesture and beat gesture, were then created. Each word was given 1 for the meaningful gesture variable if there is a corresponding meaningful gesture annotated, and was given 1 for beat gesture if it is overlapping with the stroke of a beat gesture. We checked the reliability of coding for meaningful and beat gestures, by asking a second expert coder to annotate 10% of the videos already annotated (Experiment 1: inter-rater agreement=95.3, kappa=0.922, p<. 001; Experiment 2: inter-rater agreement between coder A and C=95.6, kappa=0.929, p<.001, inter-rater agreement between coder B and C=96.7, kappa=0.948, p<.001)

#### Mouth informativeness

(Experiment 1: mean informativeness= 0.65, SD=0.28; Experiment 2: mean informativeness=0.67, SD=0.29) was quantified per word in separate online experiments. We asked an actress to produce each content word individually, and asked participants to guess the identity of the word without giving any options. Participants (recruited from Prolific) were paid £6 per hour to guess the words based on the mouth shape in an online experiment using Gorilla, with every word rated by 10 participants. Participants watched each word 2 times before typing their answer. We then calculated the averaged phonological distance between the guesses and the answer using the Python library PanPhon^80^: a distance of 0 indicates a perfect guess, and the longer the distance, the less informative the mouth movement. We then multiplied the resulting distance by −1 to create a regressor in the analysis (so that higher numbers indicate more informative mouth movement).

### Procedure

#### Experiment 1

After three passages of practice trials, each participant was presented with 50 gesture and 50 no-gesture videos in a randomized order using Presentation software (V. 18.0, www.neurobs.com). Each passage was separated by a 2000ms interval. The experiment was counterbalanced across every two participants so that they watched the videos in the same order but with counterbalanced gesture/no-gesture conditions. One-third of the videos (35) were followed by yes/no questions about the content of the video to ensure participants paid attention during the experiment and to acquire behavioural responses (See analysis in S.M.). Participants sat comfortably one meter away from the screen with a resolution of 1024*768, wearing 50Ω headphones, and were instructed to watch the videos carefully and to answer to the questions as quickly and accurately as possible (prioritizing accuracy) by pressing the left (“Yes”) or right (“No”) control key. Participants were asked to avoid moving, keep their facial muscles relaxed and reduce blinking, but they were also told that it is better to blink occasionally than to avoid blinking because of potential discomfort due to e.g., drying of the eyes. Similar instructions were written on the screen. The recording took thirty minutes with three breaks. Analyses of behavioural responses are reported in the supplementary materials.

#### Experiment 2

The procedure was almost identical to the first experiment. The only exceptions are: we used four passages as practice; we used 39/40 with gesture/without gesture videos; the interval between videos are modified to 1000ms; and 40 passages were followed by comprehension questions. The recording lasted an average of 60 minutes.

### EEG Recording

For both experiments, a 32-channel BioSimi system with silver-silver chloride electrodes and 24 bit resolution was used for the EEG recording, following a 10-10 international system layout. A common reference included the CMS electrode (serving as the online reference) and DRL electrode (serving as the ground electrode). Elastic head caps were used to keep the electrodes in place. Two external electrodes were attached under the left and right mastoids for off-line reference, while two other external eye electrodes were attached below the left eye and on the right canthus to detect blinks and eye movements. Electrolyte gel was inserted on each electrode to improve connectivity between the skull and the electrode. To check for relative impedance differences, the electrode offsets were kept between +/-25mV The recording was carried out in a shielded room with the temperature kept at 18 °C.

### EEG pre-processing

The raw data were pre-processed with EEGLAB (version 14.1.1)^81^ and ERPLAB (version 7.0.0)^73^ in MATLAB (R2017b, https://www.mathworks.com/products/matlab.html). All electrodes were included. Triggers were sent at the onset of each video, and word onset was subsequently calculated from the word boundary annotation. Any lag between trigger and stimuli presentation was also measured and corrected (Experiment 1: Mean= 210.33ms, SD=69.92; Experiment 2: Mean=40.03ms, SD=1.68). The EEG file was re-referenced to average of the left and right mastoids (M1 and M2), down-sampled from 2048Hz to 256Hz to speed up preprocessing, and separated into epochs each containing data from −100 to 1200ms around word onset^60^. The data was filtered with a second order Butterworth 0.05-100Hz band-pass filter. Due to the likely overlap between any baseline period (−100 to 0ms) and the EEG signal elicited by the previous word, we did not perform baseline correction, but instead extracted the mean EEG amplitude in this time interval and later used it as a control variable in regression analysis. This method is widely adopted in experiments using more naturalistic stimuli.^60,68^ We conducted independent component analysis based artifact correction (ICA). Two independent experts manually labelled eye movement and other noise (e.g. heart beat, line noise) components that were subsequently removed from the data. Further artifact rejection was conducted by first using a moving window peak-to-peak analysis (Voltage Threshold=100 μV, moving window full width=200 ms, window step=20 ms) and then steplike artifact analysis (Voltage Threshold=35 μV, moving window full width=400 ms, window step=10 ms). This resulted in an average rejection of 12.43% (SD=12.49) of the data in Experiment 1, and 12.18% (SD=14.43) of the data in Experiment 2. The ERP files were then computed from pre-processed data files, and were additionally filtered with a 30Hz low-pass filter.

### EEG Analysis: Hierarchical Linear Modeling (for Experiment 1)

We used the LIMO (hierarchical LInear MOdeling) toolbox^63^ working under MATLAB (R2017b, https://www.mathworks.com/products/matlab.html). For each participant, we created a single-trial file from the EEG file, and a continuous variable containing surprisal of each word. In the first level analysis for each participant, a regression analysis was performed for each data point in 0-1200ms time window per electrode per word, with EEG voltage as the dependent variable and word surprisal as the independent variable, thus generating a matrix of beta values, which indicate whether and when surprisal has an effect for each participant. In the second level of the analysis across all participants, the averaged beta matrix was compared with 0 using a one-sample t-test (bootstrap set at 1000, clustering corrected against spatial and temporal multiple comparison)^83^.

### EEG Analysis: Linear Mixed Effect Regression Analysis

#### Experiment 1

After determining the time window where surprisal has an effect, we performed linear mixed effect analysis (LMER) on the resulting 300-600ms time window. The analysis was conducted using the lme4^84^ package running under R Studio (version 3.4.1, http://www.rstudio.com/). We excluded from the analyses: (a) all function words, modal words, and proper names; (b) words without a surprisal value (26 words, due to the lack of occurrence of the combination between the word and its context in the corpus); (c) words without a mean F0 score (4 words, due to insufficient data points when calculating the average); (d) words associated with both beat and meaningful gestures (3 words); (e) words occurring without any gesture in the “with gesture” condition, and the corresponding words in without gesture videos (406 words). This was done to avoid data unbalance as there were three times more words with no gestures (combining the videos with and without gestures). Mean ERP in the 300-600ms time window (as determined in the prior hierarchical linear modeling step) was extracted from 32 scalp electrodes for each word and was used as the dependent variable. Mean ERP in the −100 to 0ms time window was extracted as the baseline. The independent variables included 1) predictors: log-transformed surprisal, mean F0, meaningful gestures, beat gestures, mouth informativeness, and all up to three-way interactions between surprisal and any two cues, excluding interactions containing meaningful gesture*beat gestures (as the three instances were removed from the data), 2) control variables: baseline extracted between −100 to 0ms, word frequency, word length, word order in the sentence, sentence order in experiment, relative position of each electrode measured by its X, Y and Z coordinate position^85^ acquired from BioSemi website (https://www.biosemi.com/download.htm). No main or interaction effects showed multicollinearity, with variance inflation factor (VIF) less than 2, kappa=4.871. All continuous variables, including ERP, surprisal, mean F0, mouth informativeness, baseline, frequency, word length, word order, sentence order and X, Y, Z position of electrodes were scaled using the “scale” function in R, so that each coefficient represents the effect size of the variable. Surprisal and frequency were log-transformed to normalize the data. All categorical variables were sum coded so that each coefficient represents the size of the contrast from the given predictor value compared with the grand mean (intercept)^68^.

We further included word types and participant as random intercept in the random structure. We attempted to construct a maximal random structure by entering all main and interactions as a random slope of participants, but the model failed to converge. As a result, we included the highest interaction (three-way interactions) as random slope for participants^69^, and surprisal was included as random slope for lemma. Our analysis included 31 participants, 381 word type lemmas and 480,212 data points.

#### Experiment 2

The replication study analyzed the mean ERP within the 300-600ms time window following the same process. We included the highest interaction of within participant variables (three-way interactions between surprisal and cues) as random slope for participants, but we did not include surprisal as random slope for item due to convergence issue. No main or interaction effects showed multicollinearity, with variance inflation factor (VIF) less than 2.5, kappa=5.76. Our analysis included 20 participants, 510 word type lemmas and 434,944 data points.

## Authorship contribution statement

Vigliocco conceived the idea for the study, Frassinelli carried out the computational modelling, contributed to the design of the study and the statistical analyses, Tuomainen contributed to the design of the study and the analysis. Zhang, designed, conducted, analysed and interpreted the EEG data. Zhang, Vigliocco and Skipper wrote the manuscript. All authors read and approved the submitted version of the manuscript.

## Acknowledgements

The work reported here was supported by a European Research Council Advanced Grant (ECOLANG, 743035) and Royal Society Wolfson Research Merit Award (WRM\R3\170016) to GV We acknowledge the Institute for Natural Language Processing (University of Stuttgart) for the usage of their server infrastructure. The authors declare no competing financial interests.

