## Supplemental Materials for "Word predictability, prosody, gesture and mouth movements in face-to-face language comprehension"

***More than words: Word predictability, prosody, gesture and mouth movements in natural language comprehension***

1. Results: full results and EEG plots

*Table 1.*

Full result: linear mixed effects regression model on N400 (300-600ms).

| | Experiment 1 | | | | Experiment 2 | | | |
|---|---|---|---|---|---|---|---|---|
| Fixed Effects | β | Std Error | t | p | β | Std Error | t | p |
| (Intercept) | 0.007 | 0.010 | 0.732 | 0.466 | 0.003 | 0.009 | 0.327 | 0.745 |
| Predictor Variables | | | | | | | | |
| Surprisal | -0.007 | 0.014 | -0.502 | 0.616 | -0.067 | 0.004 | -18.094 | < .001*** |
| Mean F0 | 0.011 | 0.002 | 5.262 | < .001*** | 0.014 | 0.002 | 7.625 | < .001*** |
| Mouth Informativeness | 0.013 | 0.008 | 1.652 | 0.100 | 0.010 | 0.005 | 1.907 | 0.057 |
| Meaningful Gesture (Present) | 0.006 | 0.001 | 5.232 | < .001*** | 0.007 | 0.001 | 5.853 | < .001*** |
| Beat Gesture (Present) | -0.004 | 0.001 | -2.500 | 0.012* | -0.006 | 0.001 | -4.002 | < .001*** |
| Surprisal:Mean F0 | 0.022 | 0.003 | 7.467 | < .001*** | 0.012 | 0.002 | 5.948 | < .001*** |
| Surprisal:Mouth Informativeness | 0.018 | 0.015 | 1.221 | 0.224 | -0.013 | 0.004 | -3.312 | 0.001** |
| Surprisal:Meaningful Gesture (Present) | 0.007 | 0.001 | 4.653 | < .001*** | 0.011 | 0.001 | 8.601 | < .001*** |
| Surprisal:Beat Gesture (Present) | -0.009 | 0.002 | -5.345 | < .001*** | -0.010 | 0.001 | -7.125 | < .001*** |
| Mean F0:Mouth Informativeness | -0.002 | 0.002 | -1.553 | 0.120 | 0.000 | 0.001 | 0.235 | 0.814 |
| Mean F0:Meaningful Gesture (Present) | 0.005 | 0.001 | 4.148 | < .001*** | 0.005 | 0.001 | 4.570 | < .001*** |
| Mean F0:Beat Gesture (Present) | -0.003 | 0.002 | -1.924 | 0.054 | 0.009 | 0.002 | 5.689 | < .001*** |

| | | | | | | | | |
|---|---|---|---|---|---|---|---|---|
| Mouth Informativeness:Meaningful Gesture (Present) | 0.004 | 0.001 | 3.072 | 0.002** | 0.007 | 0.001 | 6.210 | < .001*** |
| Mouth Informativeness:Beat Gesture (Present) | 0.012 | 0.002 | 8.005 | < .001*** | 0.004 | 0.001 | 3.181 | 0.001*** |
| Surprisal:Mean F0:Mouth Informativeness | 0.008 | 0.007 | 1.164 | 0.252 | -0.008 | 0.004 | -2.151 | 0.042* |
| Surprisal:Mean F0:Meaningful Gesture (Present) | 0.001 | 0.006 | 0.094 | 0.926 | -0.004 | 0.003 | -1.023 | 0.318 |
| Surprisal:Mean F0:Beat Gesture (Present) | 0.004 | 0.006 | 0.581 | 0.565 | 0.007 | 0.005 | 1.377 | 0.182 |
| Surprisal:Mouth Informativeness:Meaningful Gesture (Present) | -0.016 | 0.006 | -2.830 | 0.008** | -0.005 | 0.004 | -1.190 | 0.247 |
| Surprisal:Mouth Informativeness:Beat Gesture (Present) | 0.003 | 0.004 | 0.642 | 0.525 | 0.003 | 0.004 | 0.726 | 0.475 |
| Control Variables | | | | | | | | |
| Word Order | -0.011 | 0.002 | -4.778 | < .001*** | 0.000 | 0.002 | -0.241 | 0.809 |
| Word Length | -0.013 | 0.004 | -2.847 | 0.004** | -0.004 | 0.003 | -1.456 | 0.145 |
| Sentence Order | -0.007 | 0.001 | -7.868 | < .001*** | 0.001 | 0.001 | 1.478 | 0.139 |
| Baseline | 0.788 | 0.001 | 862.209 | < .001*** | 0.803 | 0.001 | 884.984 | < .001*** |
| Electrode X | -0.006 | 0.001 | -6.491 | < .001*** | -0.007 | 0.001 | -7.537 | < .001*** |
| Electrode Y | 0.008 | 0.001 | 8.387 | < .001*** | 0.010 | 0.001 | 10.674 | < .001*** |
| Electrode Z | 0.001 | 0.001 | 1.271 | 0.204 | -0.005 | 0.001 | -5.909 | < .001*** |

| | Experiment 1 | Experiment 2 |
|---|---|---|

| Random Effects | | Variance | Std.Dev. | Variance | Std.Dev. |
|---|---|---|---|---|---|
| Lemma | (Intercept) | 0.012 | 0.108 | 0.014 | 0.118 |
| | Surprisal | 0.044 | 0.211 | - | - |
| Participant ID | (Intercept) | 0.001 | 0.034 | 0.001 | 0.031 |
| | Surprisal:Mean F0:Mouth Informativeness | 0.001 | 0.039 | 0.000 | 0.016 |
| | Surprisal:Mean F0:Meaningful Gesture (Present) | 0.001 | 0.030 | 0.000 | 0.015 |
| | Surprisal:Mean F0:Beat Gesture (Present) | 0.001 | 0.033 | 0.000 | 0.021 |
| | Surprisal:Mouth Informativeness:Meaningful Gesture (Present) | 0.001 | 0.030 | 0.000 | 0.017 |
| | Surprisal:Mouth Informativeness:Beat Gesture (Present) | 0.000 | 0.021 | 0.000 | 0.018 |

| Model: Experiment 1 | | | | |
|---|---|---|---|---|
| AIC | BIC | logLik | deviance | df.resid |
| 892159 | 892746 | -446026 | 892053 | 480159 |
| **Model: Experiment 2** | | | | |
| AIC | BIC | logLik | deviance | df.resid |
| 776081 | 776630 | -387990 | 775981 | 434894 |

*Notes.* * $p < .05$, ** $p < .01$, *** $p < .001$

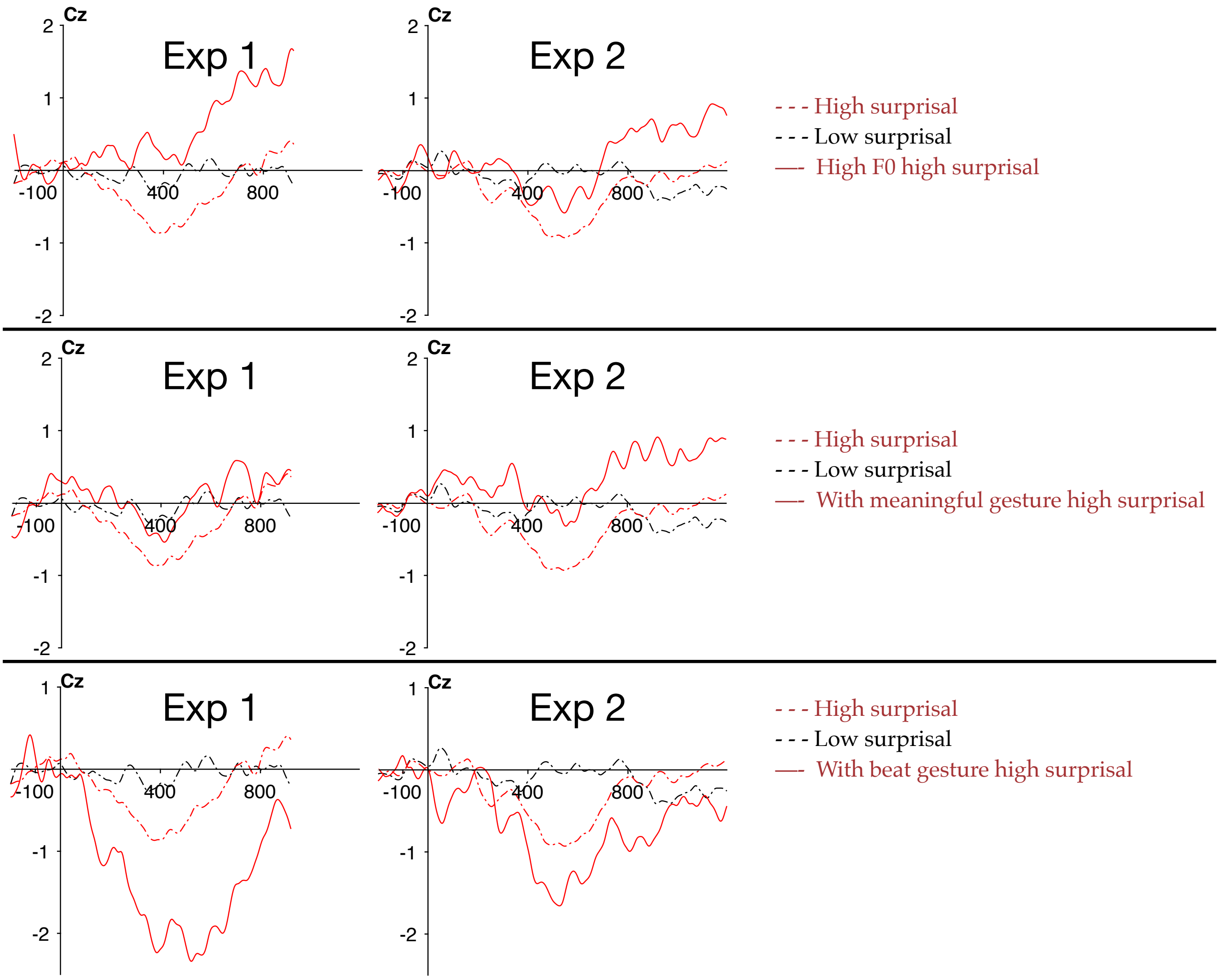


*Figure 1.*

Surprisal interact with multimodal cues in both experiments. For illustration only, continuous variables (surprisal, F0 and mouth informativeness) were categorized into high and low groups, each containing words with 30% highest/lowest values. ERP data was additionally filtered with 15Hz low pass filter for illustration. The red/black dotted lines illustrate the averaged ERP of words in high/low surprisal group, indicating the main effect of surprisal. The red solid line illustrates the averaged ERP of words with high surprisal but also with the presence of a multimodal cue, indicating that the effect of surprisal is modulated by each cue.

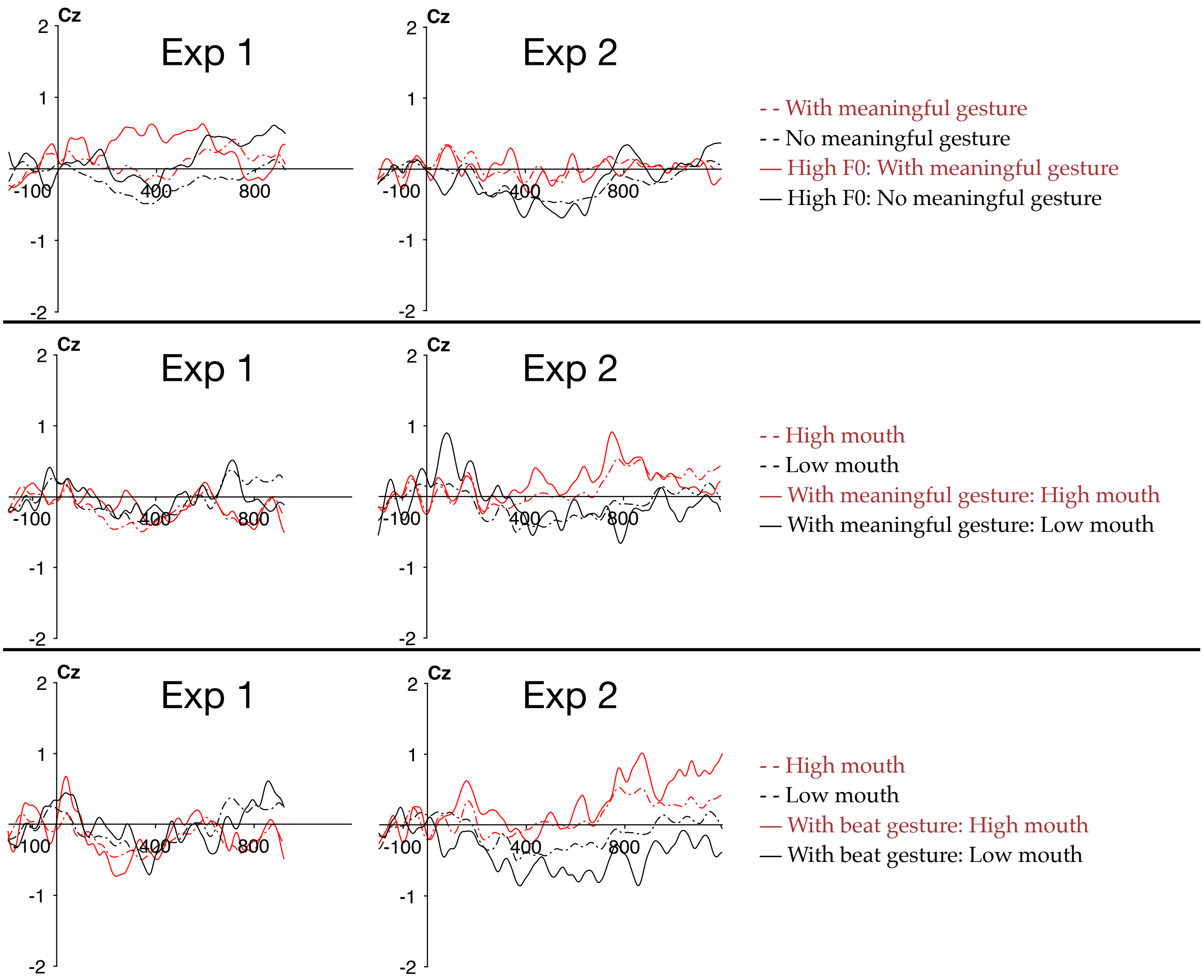


*Figure 2.*

The effect of multimodal cues interact in N400 time window. Data was categorized and additionally filtered for illustration only. Dotted lines indicate the main effect of each cue, and the solid lines indicate the effect of a cue with the presence of another cue. The effect of meaningful gesture is enlarged with high F0; similarly, the effect of mouth is enlarged with the presence of gestures.

2. Establishing the effect of surprisal

Previous study found that surprisal derived by an n-gram model can predict the N400 amplitude per word in written (Frank et al., 2015) and audio stimuli (Alday et al., 2017). However, as the audiovisual stimuli we use contains more information sources, we performed a series of analysis to test whether surprisal remains to predict language comprehension.

1) Behavioural effect of surprisal

In Experiment 1, we analysed whether averaged surprisal per passage affect the accuracy and response time for the 35 comprehension questions in order to examine the behavioural impact of surprisal. We constructed separate LMER models for accuracy and response time respectively (binomial regression model for accuracy and linear regression model for response time). We included mean surprisal per passage (calculated by averaging surprisal of all content word) as predictor variable, and participant and passageID as random intercept to control for by participant and by passage variation. All continuous variables (response time and mean surprisal) was standardised using the "scale" function, while all categorical variables (accuracy, participant and passageID) are sum coded.

We found that accuracy decreased with an increase in surprisal (Mean=82.1%, SD=0.384, $\beta$=-0.784, $p<.001$). Similarly, we found that sentences with higher averaged surprisal had slower reaction times (Mean=4129.8 ms, SD=2881.3, $\beta$=0.089, $p$=.024). The reaction time are overall longer (averaged around 4 seconds) because participants were instructed to prioritise accuracy. These findings confirm that sentences with higher surprisal were harder to process.

2) Time-window of surprisal

As is reported in the main text, we conducted hierarchical linear modelling in Experiment 1 in order to determine the EEG time-window sensitive to surprisal. We repeated the same process in Experiment 2 and found a component with a similar spatial distribution but a slightly more extended temporal distribution (around 350-850 ms post stimulus, see Figure 3 below). The longer time window may resulted from the longer duration of each word in Experiment 2 (Experiment 1: mean=440ms; Experiment 2: mean=508ms), which cause slightly longer processing time. Alternatively, this may also be related with the fact that there are more words in Experiment 2 (included lemma=510) compared with Experiment 1 (included lemma=381). This increase of word number may cause more variable processing time, causing the averaged EEG response to cover a longer time window. We repeated the same LMER analysis on ERP within 350-850ms in Experiment 2 (see table 2 below), and the results are the same with what we reported in the main text (using 300-600ms data). As a result, we used 300-600ms time window for all following analysis.

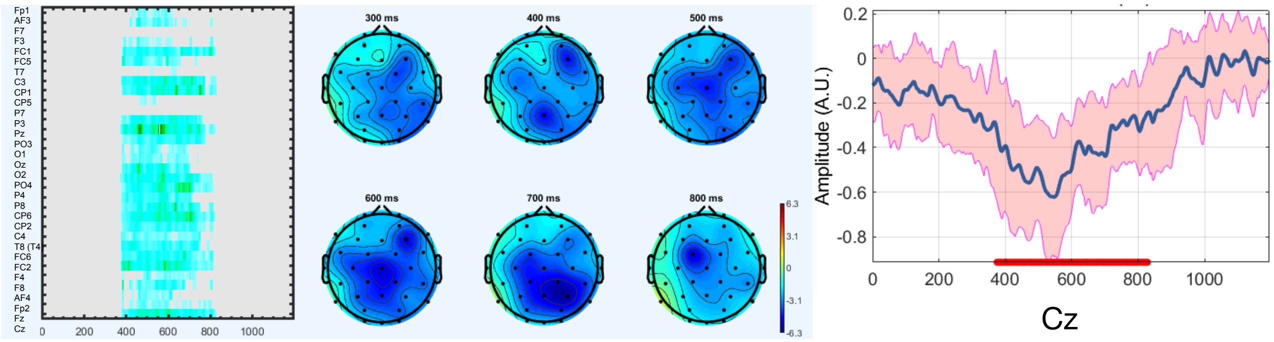


*Figure 3.*

Hierarchical linear modelling: surprisal induced a more negative ERP in 350-800ms time-window in Experiment 2.

*Table 2.*

Linear mixed effects regression model with N400 (350-850ms) as dependent variable in Experiment 2.

| Fixed Effects | β | Std Error | t | p |
|---|---|---|---|---|
| (Intercept) | -0.001 | 0.010 | -0.079 | 0.937 |
| Predictor Variables | | | | |
| Surprisal | -0.069 | 0.004 | -17.184 | <.001*** |
| Mean F0 | 0.014 | 0.002 | 7.129 | <.001*** |
| Mouth Informativeness | 0.014 | 0.006 | 2.306 | 0.022* |
| Meaningful Gesture (Present) | 0.005 | 0.001 | 4.309 | <.001*** |
| Beat Gesture (Present) | -0.009 | 0.001 | -6.358 | <.001*** |
| Surprisal:Mean F0 | 0.009 | 0.002 | 4.234 | <.001*** |
| Surprisal:Mouth Informativeness | -0.010 | 0.004 | -2.315 | 0.021* |
| Surprisal:Meaningful Gesture (Present) | 0.009 | 0.001 | 6.953 | <.001*** |
| Surprisal:Beat Gesture (Present) | -0.009 | 0.002 | -5.593 | <.001*** |
| Mean F0:Mouth Informativeness | 0.000 | 0.002 | 0.002 | 0.998 |
| Mean F0:Meaningful Gesture (Present) | 0.003 | 0.001 | 2.498 | 0.013* |
| Mean F0:Beat Gesture (Present) | 0.008 | 0.002 | 4.861 | <.001*** |
| Mouth Informativeness:Meaningful Gesture (Present) | 0.009 | 0.001 | 6.894 | <.001*** |
| Mouth Informativeness:Beat Gesture (Present) | 0.009 | 0.001 | 6.075 | <.001*** |
| Surprisal:Mean F0:Mouth Informativeness | -0.010 | 0.004 | -2.411 | 0.024* |
| Surprisal:Mean F0:Meaningful Gesture (Present) | -0.005 | 0.004 | -1.163 | 0.258 |
| Surprisal:Mean F0:Beat Gesture (Present) | 0.005 | 0.006 | 0.849 | 0.405 |
| Surprisal:Mouth Informativeness:Meaningful Gesture (Present) | -0.005 | 0.004 | -1.496 | 0.149 |
| Surprisal:Mouth Informativeness:Beat Gesture (Present) | 0.006 | 0.005 | 1.126 | 0.272 |
| Control Variables | | | | |
| Word Order | 0.000 | 0.002 | 0.125 | 0.901 |

| | | | | | |
|---|---|---|---|---|---|
| Word Length | | -0.004 | 0.003 | -1.163 | 0.245 |
| Sentence Order | | 0.002 | 0.001 | 2.213 | 0.027* |
| Baseline | | 0.770 | 0.001 | 792.635 | <.001*** |
| Electrode X | | -0.007 | 0.001 | -7.033 | <.001*** |
| Electrode Y | | 0.004 | 0.001 | 4.310 | <.001*** |
| Electrode Z | | -0.005 | 0.001 | -4.904 | <.001*** |
| Random Effects | | | | Variance | Std.Dev. |
| Lemma | (Intercept) | | | 0.017 | 0.130 |
| Participant ID | (Intercept) | | | 0.001 | 0.034 |
| | Surprisal:Mean F0:Mouth Informativeness | | | 0.000 | 0.018 |
| | Surprisal:Mean F0:Meaningful Gesture (Present) | | | 0.000 | 0.018 |
| | Surprisal:Mean F0:Beat Gesture (Present) | | | 0.001 | 0.024 |
| | Surprisal:Mouth Informativeness:Meaningful Gesture (Present) | | | 0.000 | 0.015 |
| | Surprisal:Mouth Informativeness:Beat Gesture (Present) | | | 0.000 | 0.021 |

*Notes.* * $p < .05$, ** $p < .01$, *** $p < .001$

3) Comparing surprisal calculated from different window size

We compared the surprisal value generated based on different window size in Experiment 1. For each word, we generated surprisal with varying window size n (n=1, 2, 3, 4, 5, and all), thus taking the previous n words into account when estimating the predictability of this word. In order to determine the appropriate window size, we first conducted a set of hierarchical linear modelling analysis with surprisal calculated from different window sizes as predictor variable, and then a set of multiple regression analysis with N400 (averaged ERP within 300-600ms) as the dependent variable, different surprisal and baseline ERP as independent variables (all continuous variables are standardised). As is shown in Figure 4, the all different operationalisations of surprisal induced more negative ERP within around 300-600ms. As is shown in Table 3, all operationalisations were significantly negatively related with the N400 amplitude and generated similar statistics. Since the difference between the measures are minimal, we used n=all to generate surprisal in all our subsequent analysis, thus calculating surprisal based on all previous content words.

*Figure 4.*

Hierarchical linear modelling: surprisal generated from different window sizes all induced more negative ERP within approximately 300-600ms in Experiment 1.

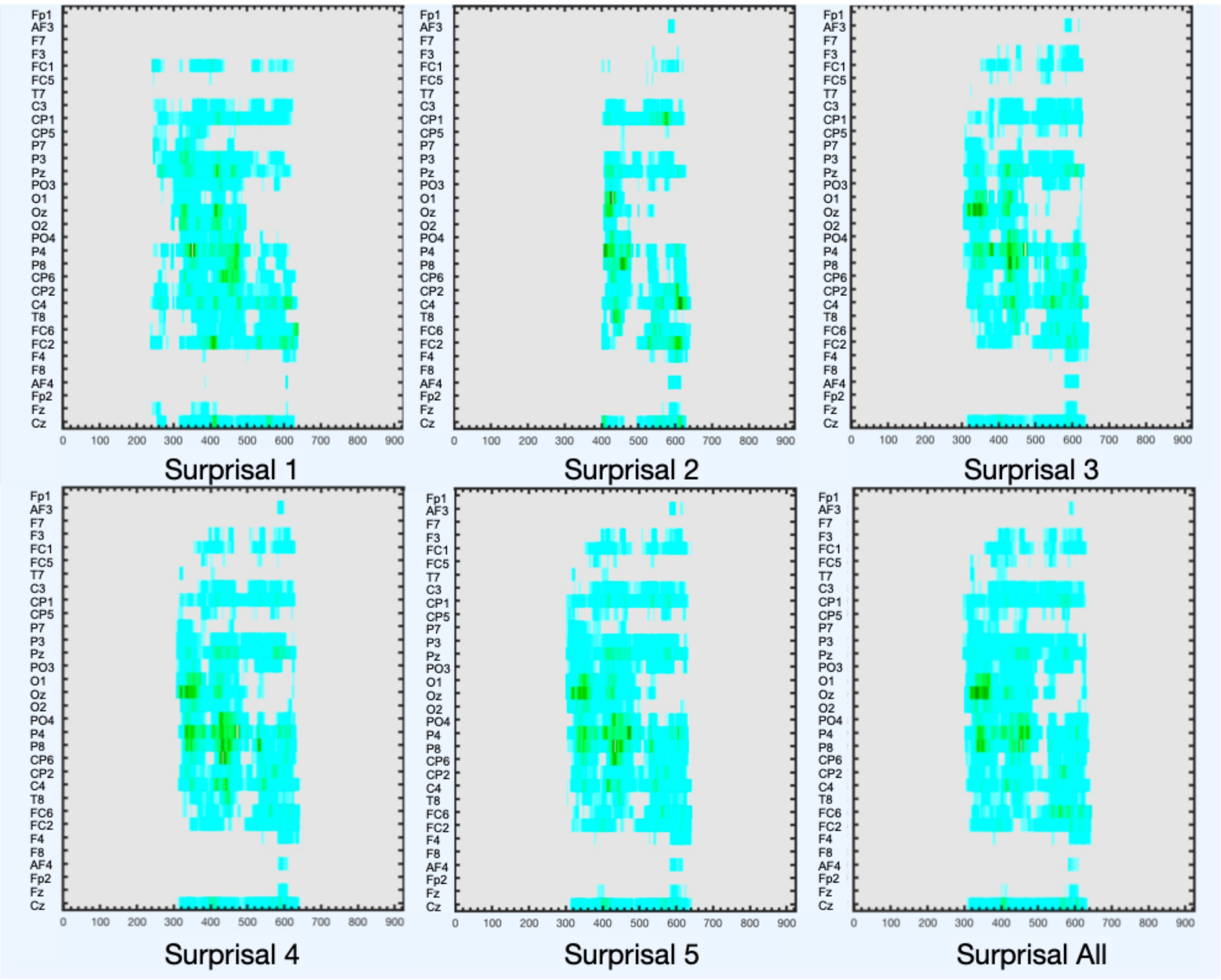


*Table 3.*

Multiple linear regression: surprisal calculated from different window sizes has similar effect on N400 (300-600ms) amplitude in Experiment 1.

| | β | Std Error | T | P | $R^2$ |
|---|---|---|---|---|---|
| Surprisal 1 | -0.008 | 0.001 | -8.479 | <.001*** | 0.613 |
| Surprisal 2 | -0.008 | 0.001 | -9.150 | <.001*** | 0.613 |
| Surprisal 3 | -0.010 | 0.001 | -11.080 | <.001*** | 0.613 |
| Surprisal 4 | -0.009 | 0.001 | -10.150 | <.001*** | 0.613 |
| Surprisal 5 | -0.009 | 0.001 | -10.050 | <.001*** | 0.613 |
| Surprisal All | -0.009 | 0.001 | -9.901 | <.001*** | 0.613 |

*Notes.* * $p < .05$, ** $p < .01$, *** $p < .001$

2. Comparison: operationalisations of prosody

Apart from mean F0, other acoustic properties (e.g. minimum F0, maximum F0, mean intensity and F0 change) may also represent prosodic accentuation. Therefore, in Experiment 1, we also compared the LMER result of these different operationalisations. We ran separate LMER model with each of the above variable as predictor, while keeping other multimodal cues constant. The results indicated that different operationalisations yielded very similar results (See table 4-7 below). Therefore, we selected mean F0, which is commonly used to represent prosody, in all our following analysis.

*Table 4.*

Linear mixed effects regression model with N400 (300-600ms) in Experiment 1 as dependent variable and minimum F0 as operationalisation of prosody.

| Fixed Effects | β | Std Error | t | p |
|---|---|---|---|---|
| (Intercept) | 0.006 | 0.010 | 0.561 | 0.575 |
| Predictor Variables | | | | |
| Surprisal | 0.012 | 0.016 | 0.734 | 0.463 |
| Min F0 | 0.024 | 0.002 | 13.662 | <.001*** |
| Mouth Informativeness | -0.004 | 0.003 | -1.309 | 0.191 |
| Meaningful Gesture (Present) | 0.006 | 0.001 | 5.048 | <.001*** |
| Beat Gesture (Present) | -0.005 | 0.001 | -3.488 | <.001*** |
| Surprisal:Min F0 | 0.003 | 0.002 | 1.426 | 0.154 |
| Surprisal:Mouth Informativeness | -0.003 | 0.002 | -1.344 | 0.179 |
| Surprisal:Meaningful Gesture (Present) | 0.010 | 0.001 | 7.071 | <.001*** |
| Surprisal:Beat Gesture (Present) | -0.011 | 0.001 | -8.630 | <.001*** |
| Min F0:Mouth Informativeness | 0.010 | 0.002 | 6.286 | <.001*** |
| Min F0:Meaningful Gesture (Present) | 0.002 | 0.001 | 1.303 | 0.193 |
| Min F0:Beat Gesture (Present) | 0.003 | 0.001 | 1.899 | 0.058 |
| Mouth Informativeness:Meaningful Gesture (Present) | -0.008 | 0.001 | -6.957 | <.001*** |

| | | | | |
|---|---|---|---|---|
| Mouth Informativeness:Beat Gesture (Present) | 0.004 | 0.002 | 2.736 | 0.006** |
| Surprisal:Min F0:Mouth Informativeness | -0.005 | 0.005 | -1.077 | 0.281 |
| Surprisal:Min F0:Meaningful Gesture (Present) | -0.006 | 0.005 | -1.200 | 0.230 |
| Surprisal:Min F0:Beat Gesture (Present) | -0.003 | 0.005 | -0.536 | 0.592 |
| Surprisal:Mouth Informativeness:Meaningful Gesture (Present) | -0.010 | 0.005 | -1.941 | 0.052 |
| Surprisal:Mouth Informativeness:Beat Gesture (Present) | 0.005 | 0.005 | 1.010 | 0.313 |
| Control Variables | | | | |
| Word Order | -0.004 | 0.002 | -2.049 | 0.040 |
| Word Length | -0.008 | 0.005 | -1.688 | 0.091 |
| Sentence Order | -0.008 | 0.001 | -8.616 | <.001*** |
| Baseline | 0.787 | 0.001 | 875.74 | <.001*** |
| Frequency | 0.032 | 0.010 | 3.247 | 0.001** |
| Electrode X | -0.006 | 0.001 | -7.070 | <.001*** |
| Electrode Y | 0.008 | 0.001 | 8.620 | <.001*** |
| Electrode Z | 0.001 | 0.001 | 0.931 | 0.352 |

| Random Effects | | Variance | Std.Dev. |
|---|---|---|---|
| Lemma | (Intercept) | 0.014 | 0.119 |
| | Surprisal | 0.049 | 0.221 |
| Participant ID | (Intercept) | 0.001 | 0.034 |
| | Surprisal:Min F0:Mouth Informativeness | 0.001 | 0.026 |
| | Surprisal:Min F0:Meaningful Gesture (Present) | 0.001 | 0.026 |
| | Surprisal:Min F0:Beat Gesture (Present) | 0.001 | 0.025 |
| | Surprisal:Mouth Informativeness:Meaningful Gesture (Present) | 0.001 | 0.027 |
| | Surprisal:Mouth Informativeness:Beat Gesture (Present) | 0.001 | 0.027 |

*Notes.* * $p < .05$, ** $p < .01$, *** $p < .001$

*Table 5.*

Linear mixed effects regression model with N400 (300-600ms) in Experiment 1 as dependent variable and maximum F0 as operationalisation of prosody.

| Fixed Effects | β | Std Error | t | p |
|---|---|---|---|---|
| (Intercept) | 0.005 | 0.010 | 0.493 | 0.622 |
| Predictor Variables | | | | |
| Surprisal | 0.010 | 0.016 | 0.633 | 0.527 |
| Max F0 | 0.006 | 0.002 | 3.183 | 0.001** |
| Mouth Informativeness | -0.003 | 0.003 | -1.044 | 0.297 |
| Meaningful Gesture (Present) | 0.005 | 0.001 | 4.589 | <.001*** |
| Beat Gesture (Present) | -0.004 | 0.001 | -2.790 | 0.005** |
| Surprisal:Max F0 | 0.010 | 0.002 | 5.081 | <.001*** |
| Surprisal:Mouth Informativeness | -0.003 | 0.002 | -1.191 | 0.234 |
| Surprisal:Meaningful Gesture (Present) | 0.007 | 0.001 | 4.775 | <.001*** |
| Surprisal:Beat Gesture (Present) | -0.012 | 0.001 | -9.404 | <.001*** |
| Max F0:Mouth Informativeness | 0.007 | 0.002 | 4.413 | <.001*** |
| Max F0:Meaningful Gesture (Present) | 0.004 | 0.001 | 3.130 | 0.002** |
| Max F0:Beat Gesture (Present) | -0.004 | 0.002 | -2.317 | 0.020* |
| Mouth Informativeness:Meaningful Gesture (Present) | -0.006 | 0.001 | -5.810 | <.001*** |
| Mouth Informativeness:Beat Gesture (Present) | 0.004 | 0.002 | 2.238 | 0.025* |
| Surprisal:Max F0:Mouth Informativeness | -0.006 | 0.004 | -1.518 | 0.129 |
| Surprisal:Max F0:Meaningful Gesture (Present) | 0.004 | 0.004 | 0.844 | 0.399 |
| Surprisal:Max F0:Beat Gesture (Present) | -0.001 | 0.005 | -0.214 | 0.831 |
| Surprisal:Mouth Informativeness:Meaningful Gesture (Present) | -0.011 | 0.005 | -2.381 | 0.017* |
| Surprisal:Mouth Informativeness:Beat Gesture (Present) | 0.002 | 0.004 | 0.375 | 0.708 |
| Control Variables | | | | |
| Word Order | -0.011 | 0.002 | -4.906 | <.001*** |

| | | | | |
|---|---|---|---|---|
| Word Length | -0.013 | 0.004 | -2.822 | 0.005** |
| Sentence Order | -0.008 | 0.001 | -8.595 | <.001*** |
| Baseline | 0.787 | 0.001 | 875.343 | <.001*** |
| Frequency | 0.035 | 0.009 | 3.717 | <.001*** |
| Electrode X | -0.006 | 0.001 | -7.067 | <.001*** |
| Electrode Y | 0.008 | 0.001 | 8.614 | <.001*** |
| Electrode Z | 0.001 | 0.001 | 0.931 | 0.352 |

| Random Effects | | Variance | Std.Dev. |
|---|---|---|---|
| Lemma | (Intercept) | 0.013 | 0.112 |
| | Surprisal | 0.046 | 0.215 |
| Participant ID | (Intercept) | 0.001 | 0.034 |
| | Surprisal:Max F0:Mouth Informativeness | 0.000 | 0.021 |
| | Surprisal:Max F0:Meaningful Gesture (Present) | 0.000 | 0.022 |
| | Surprisal:Max F0:Beat Gesture (Present) | 0.001 | 0.025 |
| | Surprisal:Mouth Informativeness:Meaningful Gesture (Present) | 0.001 | 0.024 |
| | Surprisal:Mouth Informativeness:Beat Gesture (Present) | 0.001 | 0.023 |

*Notes.* * $p < .05$, ** $p < .01$, *** $p < .001$

*Table 6.*

Linear mixed effects regression model with N400 (300-600ms) in Experiment 1 as dependent variable and mean intensity as operationalisation of prosody.

| Fixed Effects | β | Std Error | t | p |
|---|---|---|---|---|
| (Intercept) | 0.006 | 0.010 | 0.593 | 0.553 |
| Predictor Variables | | | | |
| Surprisal | 0.012 | 0.016 | 0.758 | 0.449 |

| | | | | |
|---|---|---|---|---|
| Mean Intensity | 0.004 | 0.002 | 1.920 | 0.055 |
| Mouth Informativeness | -0.008 | 0.003 | -2.660 | 0.008** |
| Meaningful Gesture (Present) | 0.007 | 0.001 | 5.766 | <.001*** |
| Beat Gesture (Present) | -0.004 | 0.002 | -2.577 | 0.010* |
| Surprisal:Mean Intensity | 0.008 | 0.002 | 4.449 | <.001*** |
| Surprisal:Mouth Informativeness | 0.004 | 0.003 | 1.743 | 0.081 |
| Surprisal:Meaningful Gesture (Present) | 0.007 | 0.002 | 4.314 | <.001*** |
| Surprisal:Beat Gesture (Present) | -0.013 | 0.001 | -10.531 | <.001*** |
| Mean Intensity:Mouth Informativeness | 0.007 | 0.002 | 4.004 | <.001*** |
| Mean Intensity:Meaningful Gesture (Present) | 0.007 | 0.001 | 6.132 | <.001*** |
| Mean Intensity:Beat Gesture (Present) | 0.004 | 0.002 | 2.669 | 0.008** |
| Mouth Informativeness:Meaningful Gesture (Present) | -0.009 | 0.001 | -7.707 | <.001*** |
| Mouth Informativeness:Beat Gesture (Present) | 0.002 | 0.002 | 1.337 | 0.181 |
| Surprisal:Mean Intensity:Mouth Informativeness | -0.007 | 0.004 | -1.753 | 0.080 |
| Surprisal:Mean Intensity:Meaningful Gesture (Present) | -0.001 | 0.005 | -0.194 | 0.846 |
| Surprisal:Mean Intensity:Beat Gesture (Present) | -0.005 | 0.005 | -0.976 | 0.329 |
| Surprisal:Mouth Informativeness:Meaningful Gesture (Present) | -0.007 | 0.004 | -1.683 | 0.092 |
| Surprisal:Mouth Informativeness:Beat Gesture (Present) | 0.004 | 0.005 | 0.861 | 0.389 |
| Control Variables | | | | |
| Word Order | -0.011 | 0.002 | -5.064 | <.001*** |
| Word Length | -0.013 | 0.004 | -2.893 | 0.004** |
| Sentence Order | -0.009 | 0.001 | -9.388 | <.001*** |
| Baseline | 0.788 | 0.001 | 875.48 | <.001*** |
| Frequency | 0.041 | 0.010 | 4.222 | <.001*** |
| Electrode X | -0.006 | 0.001 | -7.071 | <.001*** |
| Electrode Y | 0.008 | 0.001 | 8.618 | <.001*** |
| Electrode Z | 0.001 | 0.001 | 0.932 | 0.351 |

| Random Effects | | Variance | Std.Dev. |
|---|---|---|---|
| Lemma | (Intercept) | 0.013 | 0.114 |
| | Surprisal | 0.049 | 0.221 |
| Participant ID | (Intercept) | 0.001 | 0.035 |
| | Surprisal:Mean Intensity:Mouth Informativeness | 0.000 | 0.022 |
| | Surprisal:Mean Intensity:Meaningful Gesture (Present) | 0.001 | 0.027 |
| | Surprisal:Mean Intensity:Beat Gesture (Present) | 0.001 | 0.027 |
| | Surprisal:Mouth Informativeness:Meaningful Gesture (Present) | 0.001 | 0.023 |
| | Surprisal:Mouth Informativeness:Beat Gesture (Present) | 0.001 | 0.024 |

*Notes.* * $p < .05$, ** $p < .01$, *** $p < .001$

*Table 7.*

Linear mixed effects regression model with N400 (300-600ms) in Experiment 1 as dependent variable and change of F0 within a word as operationalisation of prosody.

| Fixed Effects | β | Std Error | t | p |
|---|---|---|---|---|
| (Intercept) | 0.006 | 0.010 | 0.637 | 0.524 |
| Predictor Variables | | | | |
| Surprisal | 0.014 | 0.016 | 0.884 | 0.376 |
| F0 Change | -0.007 | 0.002 | -3.450 | 0.001** |
| Mouth Informativeness | -0.002 | 0.003 | -0.632 | 0.528 |
| Meaningful Gesture (Present) | 0.006 | 0.001 | 5.239 | <.001*** |
| Beat Gesture (Present) | -0.003 | 0.001 | -2.213 | 0.027* |
| Surprisal:F0 Change | 0.004 | 0.002 | 2.242 | 0.025* |
| Surprisal:Mouth Informativeness | -0.001 | 0.002 | -0.593 | 0.553 |
| Surprisal:Meaningful Gesture (Present) | 0.007 | 0.001 | 5.024 | <.001*** |

| | | | | |
|---|---|---|---|---|
| Surprisal:Beat Gesture (Present) | -0.011 | 0.001 | -8.542 | <.001*** |
| F0 Change:Mouth Informativeness | 0.004 | 0.002 | 2.437 | 0.015* |
| F0 Change:Meaningful Gesture (Present) | 0.001 | 0.001 | 0.952 | 0.341 |
| F0 Change:Beat Gesture (Present) | -0.005 | 0.002 | -3.467 | 0.001** |
| Mouth Informativeness:Meaningful Gesture (Present) | -0.007 | 0.001 | -6.034 | <.001*** |
| Mouth Informativeness:Beat Gesture (Present) | 0.005 | 0.002 | 2.835 | 0.005** |
| Surprisal:F0 Change:Mouth Informativeness | -0.004 | 0.004 | -0.874 | 0.382 |
| Surprisal:F0 Change:Meaningful Gesture (Present) | 0.002 | 0.004 | 0.553 | 0.580 |
| Surprisal:F0 Change:Beat Gesture (Present) | -0.001 | 0.004 | -0.338 | 0.735 |
| Surprisal:Mouth Informativeness:Meaningful Gesture (Present) | -0.010 | 0.005 | -2.173 | 0.030* |
| Surprisal:Mouth Informativeness:Beat Gesture (Present) | 0.003 | 0.005 | 0.535 | 0.592 |
| Control Variables | | | | |
| Word Order | -0.009 | 0.002 | -4.073 | <.001*** |
| Word Length | -0.010 | 0.004 | -2.341 | 0.019* |
| Sentence Order | -0.008 | 0.001 | -8.950 | <.001*** |
| Baseline | 0.787 | 0.001 | 875.02 | <.001*** |
| Frequency | 0.037 | 0.009 | 3.925 | <.001*** |
| Electrode X | -0.006 | 0.001 | -7.064 | <.001*** |
| Electrode Y | 0.008 | 0.001 | 8.614 | <.001*** |
| Electrode Z | 0.001 | 0.001 | 0.929 | 0.353 |

| Random Effects | | Variance | Std.Dev. |
|---|---|---|---|
| Lemma | (Intercept) | 0.012 | 0.111 |
| | Surprisal | 0.046 | 0.215 |
| Participant ID | (Intercept) | 0.001 | 0.034 |
| | Surprisal:F0 Change:Mouth Informativeness | 0.001 | 0.023 |
| | Surprisal:F0 Change:Meaningful Gesture (Present) | 0.000 | 0.021 |
| | Surprisal:F0 Change:Beat Gesture (Present) | 0.001 | 0.023 |

| | | | |
|---|---|---|---|
| | Surprisal:Mouth Informativeness:Meaningful Gesture (Present) | 0.001 | 0.025 |
| | Surprisal:Mouth Informativeness:Beat Gesture (Present) | 0.001 | 0.025 |

*Notes*. * $p < .05$, ** $p < .01$, *** $p < .001$
